# Modeling the evolutionary architectures of human enhancer sequences reveals distinct origins, functions, and associations with human-trait variation

**DOI:** 10.1101/2020.08.03.235051

**Authors:** Sarah L. Fong, John A. Capra

**Affiliations:** Vanderbilt Genetics Institute, Vanderbilt University, Nashville, TN, USA; Department of Biological Sciences, Vanderbilt University, Nashville TN, USA; Bakar Computational Health Sciences Institute and Department of Epidemiology and Biostatistics, University of California, San Francisco, USA

## Abstract

**Motivation:** Despite the importance of gene regulatory enhancers in human biology and evolution, we lack a comprehensive evolutionary model of enhancer sequence architecture and function. This substantially limits our understanding of the genetic basis for divergence between species and our ability to interpret the effects of non-coding variants on human traits.

**Results:** To explore enhancer sequence evolution and its relationship to regulatory function, we traced the evolutionary origins of human sequences with enhancer activity defined by eRNA from diverse tissues and cellular contexts. The majority of enhancers are sequences of a single evolutionary age (“simple” enhancer architectures), likely indicating constraint against genomic rearrangements. A minority of enhancers are composites of sequences of multiple evolutionary ages (“complex” enhancer architectures). Compared to simple enhancers, complex enhancers are older, more pleiotropic, and more active across species. Genetic variants within complex enhancers are also less likely to have effects on human traits and biochemical activity. Transposable-element-derived sequences have made diverse contributions to enhancer architectures; some have nucleated enhancers with simple architectures, while others have remodeled older sequences to create complex regulatory architectures.

**Conclusions:** Based on these results, we propose a framework for modeling enhancer sequence architecture and evolution. Applying this framework to human enhancer sequences reveals multiple, distinct trajectories of human regulatory sequence evolution. Considering these evolutionary histories can aid interpretation of the effects of variants on enhancer function.

## INTRODUCTION

Enhancers are non-coding DNA sequences bound by transcription factors that regulate gene transcription and establish tissue- and cell-specific gene expression patterns (Shlyueva et al., 2014). Rapid turnover of sequences with enhancer activity is a common evolutionary process that contributes to species-specific gene regulation and phenotypic diversity (Wittkopp & Kalay, 2012). Despite the importance of gene regulatory enhancers in human biology and evolution, we lack a comprehensive model of their evolutionary and functional dynamics.

Comparative genomic studies have demonstrated that gene regulatory activity turns over rapidly between species. For example, a comparison of enhancer-associated histone modifications revealed that active liver enhancers are rarely shared among 20 placental mammals, though most liver enhancer sequences were alignable between species (Villar et al., 2015). Similar patterns have been observed in transcription factor (TF) DNA binding events. The majority of liver TF binding events among five vertebrates are private to a single species, and DNA binding site divergence between species is largely explained by lineage-specific mutations that activate and inactivate binding sites (Schmidt et al., 2010).

Despite that enhancer activity is often species-specific, most sequences underlying active enhancers are shared across species. For example, 80% of mouse DNase hypersensitive sites (DHS) originate from the last common ancestor of mice and humans, yet only 36% of DHS sites have shared activity between humans and mice (Vierstra et al., 2014). Similarly, a comparison of human, rhesus, and mouse enhancers involved in embryonic limb development showed that most human-specific gains in enhancer activity occur in ancient mammalian sequences, most often due to a small number of substitutions (Cotney et al., 2013). These studies indicate that most enhancer sequences do not maintain consistent activity over evolutionary distances and suggest that the evolution of new functions in DNA sequences with ancient origins (sometimes referred to as exaptation) has been a common mode of enhancer evolution. Thus, it is important to distinguish the evolutionary history of enhancer activity, which is often species specific, from the history of the underlying DNA sequence, which is often ancient. For brevity, we use the term “enhancer” when discussing sequence with enhancer activity in a context of interest.

Species-specific patterns of enhancer activity can arise from a range of genomic changes. Human-specific adaptive nucleotide substitutions in conserved developmental enhancers have been shown to drive robust *in vivo* reporter activity in mouse compared with chimpanzee and rhesus orthologs (Capra, et al., 2013a; Prabhakar et al., 2008). Despite this, most gains of enhancer activity are not under strong positive selection (Pollard et al., 2006). Repetitive sequences derived from transposable elements (TEs) also contribute to species-specific enhancer activity (Chuong et al., 2017). Over evolutionary time, specific TE insertions provide new TF binding motifs, expand human gene regulatory regions, and, in some cases, associate with evolutionary shifts in nearby gene expression (Marnetto et al., 2018). Though important, TE derived sequences (TEDS) are depleted for enhancer activity compared to the rest of the genome (Emera et al., 2016; Simonti et al., 2017). Together, these results illustrate that enhancer evolution is dynamic and can proceed through different evolutionary trajectories.

Determining evolutionary origins by estimating *sequence age*—i.e. the common ancestor in which a homologous sequence first appeared—has provided useful information about enhancers’ biological functions and associations with complex human diseases. Cross-species analyses of gene regulatory elements revealed three periods of regulatory sequence innovation during vertebrate evolution. Regulatory elements of different ages were shown to have different functional targets (Lowe et al., 2011). Human enhancers with older sequence ages were recently shown to be more enriched for heritability of complex traits than enhancers in younger sequences, independent of the conservation of enhancer function across species (Hujoel et al., 2019).

A pioneering analysis of human neocortical enhancers demonstrated that enhancer sequences can be composites of sequences from multiple ages and origins (Emera et al., 2016). In light of this sequence age complexity, a two-step life cycle model was proposed to explain enhancer sequence evolution. In the first step, short proto-enhancer sequences of a single evolutionary origin gain weak enhancer activity, and most of these proto-enhancers are inactivated over time. In the second step, a fraction of proto-enhancers gain more robust activity through the integration of younger sequences carrying relevant TF binding sites (TFBSs) that may create or modify TF-complex interactions and solidify enhancer functions.

Thus, previous work has established that the evolutionary histories of enhancer sequences are informative about their functions. However, previous work has largely overlooked the *evolutionary architecture* of enhancers*—*i.e. the evolutionary sequence age(s) of sequences with enhancer activity—which more precisely reflects the evolutionary events that produced them. Thus, there is a gap in our understanding of the evolutionary dynamics that produce sequences with enhancer activity and *how* these histories relate to gene regulatory function.

Here, we build on previous enhancer sequence age strategies (Emera et al., 2016; Hujoel et al., 2019; Lowe et al., 2011; Marnetto et al., 2018) to quantify enhancer *sequence age architecture—*the age of every base pair within a sequence with enhancer activity—across human enhancers. We then evaluate how sequence age architecture relates to enhancer function, evolutionary stability, and tolerance to human variation. We emphasize that this work estimates the age of sequences with enhancer activity in humans; this estimate does not identify the time when the sequence first gained enhancer activity. Across tissues, we find the majority of enhancer sequence architectures have a single age, and a minority of enhancer sequences have multiple ages. Further, we find tissue pleiotropy, cross-species activity, and evolutionary conservation are higher in enhancers with multi-age sequence architectures, while functional differences in enhancer activity due to natural human variation occurs more frequently in single-age architectures. Based on these patterns in enhancer evolutionary architectures, we present a model of enhancer sequence evolution that will provide a useful framework for dissecting the evolution and function of human enhancer sequences.

## RESULTS

### Estimating enhancer ages using vertebrate multiple species alignments

In this study, our goal is to characterize the evolutionary sequence architecture of human enhancers and its association with regulatory function. In this section, we describe the datasets and strategies used to define sequence ages and provide context necessary for interpreting our results. We analyzed human enhancers identified by two common strategies. First, our main results evaluate 30,474 autosomal enhancers identified from active enhancer RNAs (eRNAs) from the FANTOM5 consortium (Andersson et al., 2014). Second, we validate our main findings with analysis of 20,172,949 autosomal enhancers identified from histone-modified chromatin immunoprecipitation sequencing (ChIP-seq) from the Roadmap Epigenomics Mapping Consortium (Roadmap Consortium 2015), which are reported in Supplementary Material, unless otherwise noted.

We assigned sequence ages to enhancers based on the evolutionary histories of the overlapping syntenic blocks from the UCSC 46-way alignment of diverse vertebrate species spanning 600 million years of evolution (Figure 1A; Methods). For simplicity, we grouped most recent common ancestor (MRCA) nodes into 10 age categories and report sequence age as the oldest ancestral branch on which the sequence first appeared (Methods).

**Figure 1.**
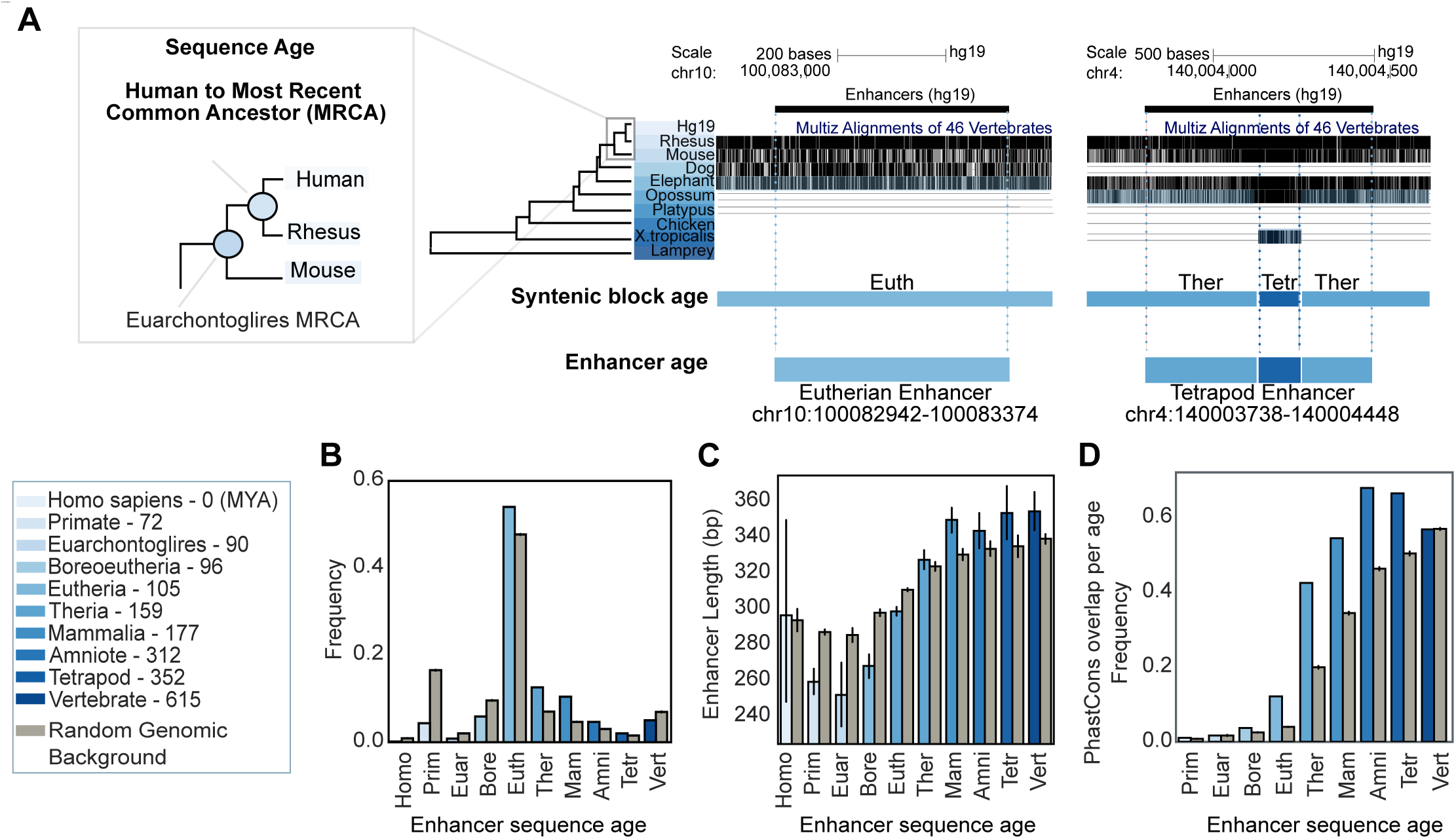
Enhancers are older, longer, and more conserved than expected from the genomic background. (A) Illustration of the method for assigning sequence ages. We quantify the age of a sequence with human enhancer activity based on the oldest most recent common ancestor (MRCA) in overlapping syntenic blocks from the MultiZ multiple sequence alignments of 46 vertebrates (inset). Enhancer age is assigned as the oldest, overlapping syntenic block age. Millions of years ago (MYA) divergence estimates from TimeTree (Hedges et al., 2015) are annotated in parenthesis in the color key. (B) The distribution of enhancer sequence ages across 30,474 FANTOM enhancers compared to 100 sets of length-matched random genomic regions (gray). Enhancers are significantly older than expected compared to length-matched random genomic regions (*p* < 2.2e-308, Mann Whitney U test). (C) Enhancer lengths by age versus 100 sets of length-matched random genomic background sets. Older enhancers are longer than expected (median 321 bp versus 310 bp, FANTOM enhancers versus random regions older than placental mammals; *p* < 2.2e-308), younger enhancers are shorter than expected (median 277 bp versus 286 bp random regions; *p* = 3e-15). (D) Enhancers are more conserved than younger enhancers and more conserved than expected (28% FANTOM enhancers versus 12% random regions overlap a PhastCons element).

### Enhancers are older, longer, and more conserved than the genomic background

Among human enhancer sequences, 54% have origins in the eutherian ancestor, while 35% can be traced to older ancestors and 11% can trace their origins to younger ancestors (Figure 1B). Human enhancers are significantly older than length-matched sets of random sequences from across the human genome (Figure 1B; *p* < 2.2e-308, Mann Whitney U). Quantifying enrichment for each sequence age, enhancers are 1.6- to 6-fold depleted of sequences younger than the eutherian ancestor and 1.2- to 2.3-fold enriched for sequences older than the eutherian ancestor (Supplemental Figure 1.2A). The older sequence age enrichment was consistent in the Roadmap enhancers (Supplemental Figure 1.3A, B). In line with previous observations (Emera et al., 2016; Lowe et al., 2011; Marnetto et al., 2018; Villar et al., 2015), sequences with human enhancer activity are older than expected from the genomic background, suggesting that they have been maintained due to their regulatory functions.

Older enhancer sequences (origins before the eutherian ancestor) are significantly longer than younger enhancer sequences (origins from eutherian ancestor and more recent) (median 321 bp versus 277 bp; p = 8.2e-122, Mann Whitney U) and longer than expected from age-matched regions from the random genomic background sets (median 321 bp versus 310 bp; *p*< 2.2e-308) (Figure 1C, Supplemental Figure 1.2B). Younger enhancers are shorter than expected (median 277 bp versus 286 bp; *p* = 3.0e-15). Similar length trends were observed in Roadmap enhancers, though only the oldest were significantly longer than expected (Supplemental Figure 1.3C). Therefore, across ages, enhancers’ lengths vary, and these lengths are different from age-matched random regions of the genome.

As expected, older enhancers are more conserved than younger enhancers and more conserved than expected from background (Figure 1E). We find that 44–69% of older enhancer ages overlap PhastCons conserved elements, while only 1–11% of younger enhancer ages overlap PhastCons elements (Supplementary Figure 1.1C). This highlights that sequence age and conservation provide complementary information; age estimates the origin of the sequence, while conservation estimates constraint on sequence variation.

### Enhancers are enriched for simple evolutionary sequence architectures

The majority (64%) of human FANTOM enhancers are found within a single syntenic block and thus a single age (Figure 2A). We refer to these as enhancers with *simple sequence age architectures* (“simple” enhancers). A minority (36%) of human FANTOM enhancers include sequences with multiple different ages, and we refer to these as enhancers with *complex sequence age architectures* (“complex” enhancers). We classified complex enhancers by the oldest sequence age, and note that human-specific enhancers can only be classified as simple enhancers because the oldest sequence age maps to the human branch (Methods). The median enhancer length is 292 bp, and the median genome-wide syntenic block is 54 bp. Thus, it was surprising that only 36% of enhancers mapped to more than one syntenic age (Figure 2B). We compared this result to length-matched random genomic regions to evaluate whether complex enhancers were under-represented given the length distributions of enhancers and syntenic blocks.

**Figure 2.**
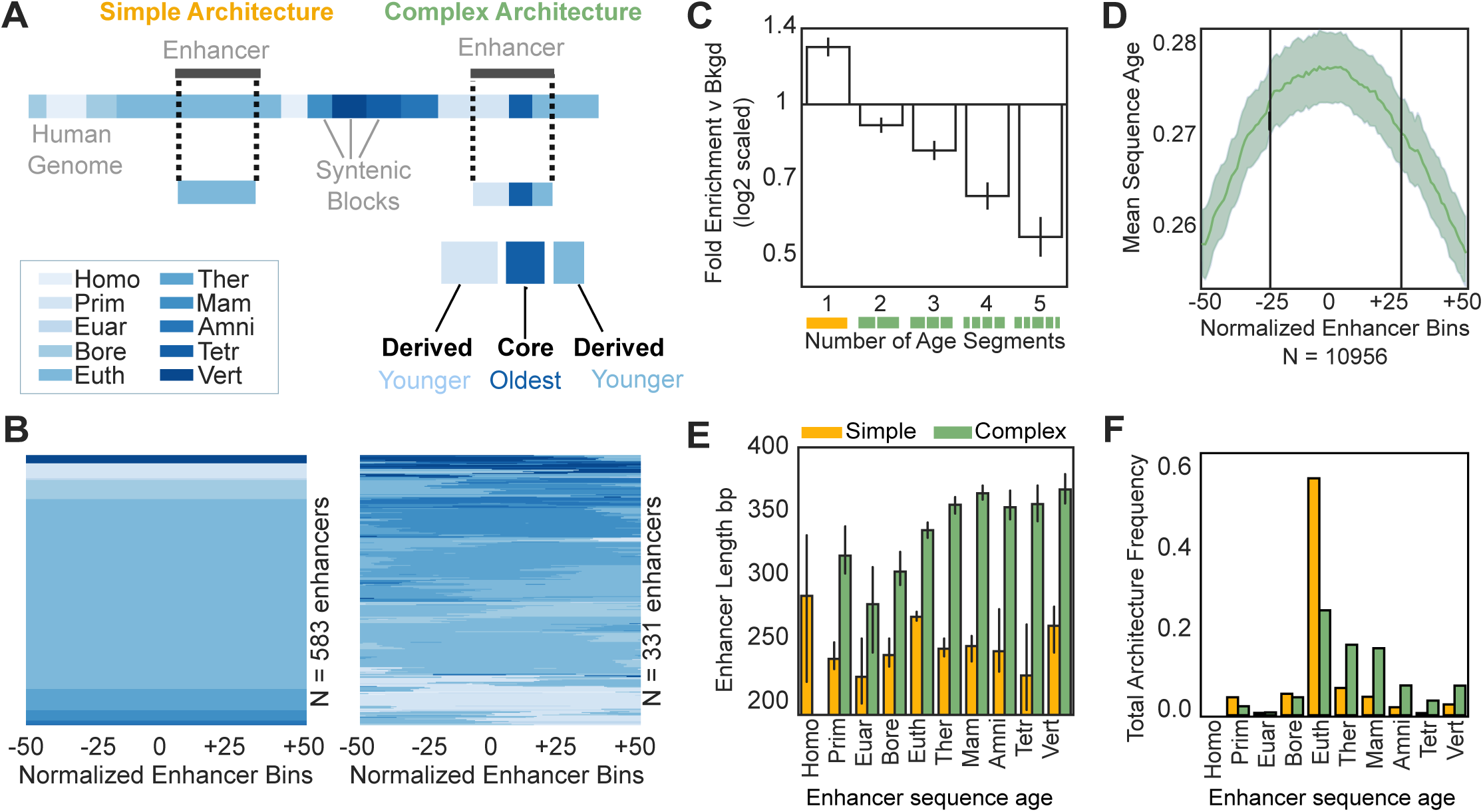
Simple and complex enhancers have distinct sequence architectures, lengths, and ages. (A) Schematic of simple and complex enhancer sequence architectures based on overlapping syntenic block ages. Simple enhancers are composed of sequence of one evolutionary age, while complex enhancers contain sequence of multiple ages. Within complex enhancers, the oldest segment is the “core” and younger segments are “derived.” (B) Example simple and complex enhancer architectures from 921 random autosomal FANTOM enhancers. The majority (64%) of FANTOM enhancers have simple architectures. For illustration, the age of each enhancer sequence is summarized across 100 equally spaced bins; the age of each sequence in each bin is indicated by color. (C) Enhancers have significantly fewer segments of different ages than expected by chance; simple FANTOM enhancers are 1.3-fold enriched versus 100 random enhancer sets (*p* = 7.6e-107, Fisher’s exact test). The fold change is calculated between the observed number of simple enhancer ages compared with the expected number of simple enhancers based on random length-matched sets. (D) Complex enhancers are on average older at their centers. Dividing complex enhancers into 100 equally spaced bins, mean complex enhancer sequence age from 10,956 complex enhancers is shown; the light green regions represent bootstrapped 95% confidence intervals. The middle 50% of complex enhancer bins are significantly older than the outer 50% of bins (0.275 versus 0.265 substitutions per site; *p* = 4.9e-166, Mann Whitney U). This pattern is mainly driven by older enhancers (Supplementary Figure 2.2). (E) Complex enhancers are longer than simple enhancers (median 347 bp versus 259 bp, *p* < 2.2e-308, error bars give 95% confidence intervals based on 10,000 bootstraps). (F) Complex enhancers are significantly older than simple enhancers (0.36 versus 0.23 mean substitutions per site, *p* < 2.2e-308).

Human enhancers are enriched for simple architectures and depleted for multiple age segments (1.3-fold enrichment simple enhancers *p* = 7.6e-107 Fisher’s exact test, 0.1–0.5-fold complex enhancers; *p* < 7.1e-12, Figure 2C, Supplemental Figure 2.4A). These differences were greatest among enhancers with therian and eutherian sequence origins (Supplemental Figure 2.4B). Roadmap enhancers are substantially longer than FANTOM enhancers (median 292 bp versus 2.4 kb), and therefore a much smaller fraction have simple architectures (median 2%, Supplemental Figure 2.9). Nonetheless, we observed similar depletion for the number of age segments in these enhancers compared with random expectation as well (Supplemental Figure 2.5A, Supplemental Figure 2.10A). This suggests constraint against insertions and deletions in sequences with gene regulatory potential.

### The oldest sequences occur in the middle of complex enhancers

Among complex enhancer sequences, we define the oldest sequence as the “core” and younger sequences as “derived” segments (Figure 2A). The core is generally at the center of the enhancer, while younger sequences are generally found on the edges of complex enhancers (Figure 2D, Methods). Stratifying enhancers by core age revealed that this pattern was driven by enhancers derived from older sequences (Supplemental Figure 2.2). In younger complex enhancers, core sequences are slightly biased towards sequence edges. This likely reflects the fact that most young complex enhancers consist of only two ages, one older and one younger (Supplemental Figure 2.4). These patterns were consistent in Roadmap enhancers, despite ChIP-peak length differences (Supplemental Figure 2.3, Supplemental Figure 2.5). This suggests that older core sequences and younger flanking sequences are non-randomly arranged within complex enhancer architectures.

### Complex enhancers are longer and older than simple enhancers

Complex enhancers are significantly longer than simple enhancers (Figure 2E and Supplemental Figure 2.5B, median 347 versus 259 bp; *p* < 2.2e-308, Mann Whitney *U* test). Some length difference is expected based on the definition of complex enhancers, since longer regions are more likely to overlap multiple syntenic blocks by chance. To evaluate whether the length difference between simple and complex enhancers was greater than expected, we shuffled length-matched enhancers and assigned architectures (simple or complex) to the resulting random regions. Repeating this 100 times, we observed that complex enhancer sequences are significantly longer than expected (median 347 bp versus 339 bp; *p* = 2.5e-06, Mann Whitney U test) and that complex enhancers have a stronger positive correlation between length and age than expected (10.7 bp/100 million years (MY), *p =* 5.2e-24 versus 5.8 bp/100 MY, *p* <2.2e-308, linear regression). In contrast, simple enhancers retain similar lengths over time (−0.8 bp/100 MY, *p* = 0.1 versus -3.5 bp/100 MY, *p* = 6.8e-296) and were slightly longer than expected (median 259 bp versus 256 bp; p = 7.3e-05, Supplemental Figure 2.6). Thus, complex enhancer sequences are longer than simple enhancer sequences, and this difference is greater than would be expected due only to the definition of complex architectures.

Next, we compared the sequence age distribution for simple and complex architectures. Sixty-eight percent of simple enhancer sequences are derived from the eutherian ancestor, while 12% are younger and 19% are older (Figure 2F and Supplemental Figure 2.5C). Simple enhancers are enriched for sequences from the eutherian ancestor, and are older than expected overall *(p <* 2.2e-308, Supplemental Figure 2.7). Conversely, 30% of complex enhancers are derived from the eutherian ancestor, 9% are younger than the eutherian ancestor and 61% of complex enhancers are older. Complex enhancers are enriched for sequences older than eutherian ancestor and are also older than expected (*p* < 2.2e-308). Though both simple and complex enhancers are enriched for older sequence ages, we observe a depletion of the oldest sequences (vertebrate ancestor) for both architectures, and simple architectures are more depleted than complex (1.4- versus 1.1-fold depleted, Supplemental Figure 2.7).

### Complex enhancers are more pleiotropic and more conserved in activity across species than simple enhancers

In this section, we evaluate whether complex and simple enhancers have differences in their patterns and breadth of activity across species. Among tissues and cell types, the ratio of active enhancers with simple versus complex sequence architectures varies. For example, the contexts with the highest fractions of simple enhancers (57–61%) include many blood cell and epithelial cell types, while the contexts with lowest fractions (49–51%) include eye, neuronal stem cell, and cardiac myocytes (Supplemental Figures 2.8, 2.9).

Enhancers with ancient origins and conserved activity across diverse mammals are known to be more pleiotropic—i.e. they have signatures of shared activity across multiple human tissues (Fish et al., 2017). Thus, we hypothesized that older complex enhancers would be more pleiotropic than younger simple enhancers. To test this, we quantified the overlap of enhancer activity across 112 tissue and cell enhancer datasets (Methods). To control for length differences between simple and complex enhancers in this and subsequent analyses, we trimmed or expanded enhancers around their midpoints to match the dataset-wide mean length (310 bp).

Complex enhancers are active across significantly more biological contexts than simple enhancers (mean of 7.4 versus 4.8 contexts; *p* = 5.9e-199, Mann Whitney U) (Figure 3A)—suggesting that complex enhancers are more pleiotropic than simple enhancers. Enhancer pleiotropy increases with age, and complex enhancers are consistently more pleiotropic than age-matched simple enhancers (Figure 3A). Analysis of trimmed Roadmap enhancers yields similar conclusions across 98 contexts; complex enhancers are more pleiotropic (Supplemental Figure 3.2). Together, these data suggest that enhancers with complex sequence age architectures are more pleiotropic and have broader function across biological contexts than enhancers with simple sequence age architectures.

**Figure 3.**
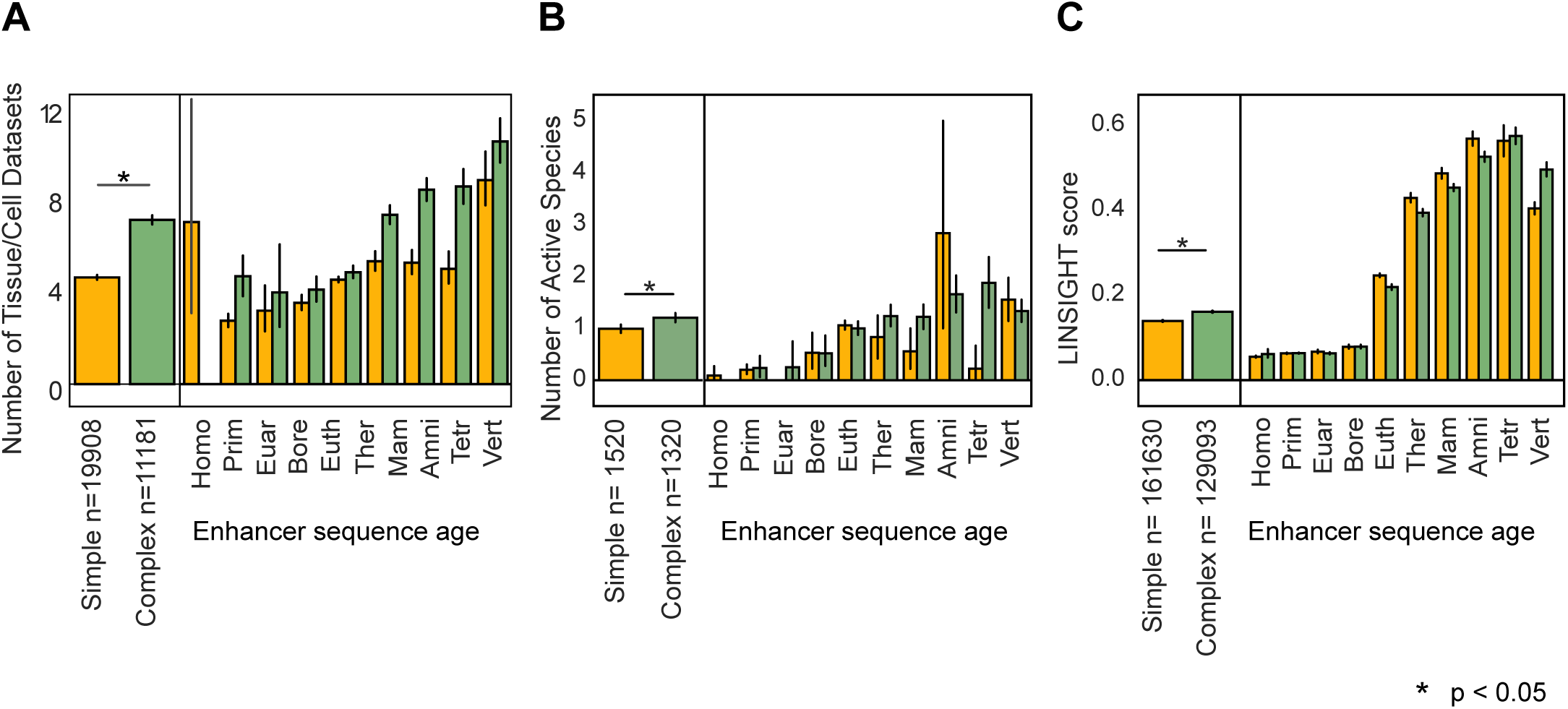
Complex enhancers are more active across tissues and species and under stronger purifying selection than simple enhancers. Simple and complex enhancer activity was evaluated across 112 FANTOM enhancer contexts. (A) Complex enhancers are more pleiotropic than simple enhancers. Overall, simple enhancers are active in 4.8 contexts on average and complex enhancers are active in 7.4 contexts (left; *p* = 5.9e-199, Mann Whitney U test). Activity across tissues increases with sequence age, but the effect is stronger for complex enhancers overall and stratified by enhancer age (right). (B) Complex human liver enhancers are active across significantly more species than simple liver enhancers (left, 1.3 versus 1.0 mean species; *p* = 4.6e-08). Simple and matched-length complex human liver enhancers defined by H3K27ac+ H3K4me3-ChIP-peaks from Villar 2015 (N =1520 and 1308) were evaluated for enhancer activity across nine placental mammals. Stratifying by enhancer age reveals that older enhancers are generally active across more species, but that the relationships between simple and complex enhancers differ in different age groups (right). (C) Complex enhancers are under stronger negative selection on the human lineage than simple enhancers (left, 0.16 versus 0.14 mean LINSIGHT score, *p* < 2.2e-308). However, estimates stratified by age generally showed similar levels among complex and simple enhancers (right). To account for length differences between architectures, all enhancers were trimmed or expanded to the mean enhancer length of 310 bp. Error bars represent 95% confidence intervals based on 10,000 bootstraps.

Given the known, positive-association between enhancer pleiotropy and cross-species activity, we next asked if simple and complex architectures differed in cross-species activity. This analysis required enhancer maps from the same tissue across species; thus, we assigned age architectures to H3K27ac+ H3K4me3-enhancers identified across liver samples from nine placental mammals (Villar et al., 2015). To control for differences in length, we randomly sampled complex enhancers matched on simple enhancer lengths (n = 1520 simple enhancers and n = 1308 matched-length complex enhancers) and evaluated cross-species activity. As expected from previous studies, human enhancers are largely species-specific, but complex liver enhancers are active across significantly more species than simple liver enhancers (1.3 versus 1.0 mean species; *p* = 4.6e-8, Mann Whitney U) (Figure 3B). In general, older enhancers are more active across species. However, we did not observe consistent activity differences between age-matched simple and complex architectures. This suggests that overall, complex enhancers are active across more species than simple enhancers and this trend is likely due to the older sequence ages of complex enhancers.

### Complex enhancers are under stronger negative selection than simple enhancers

Given the older ages, greater pleiotropy, and greater cross-species activity observed in complex enhancers, we hypothesized that they would be under stronger negative selection than simple enhancers. To evaluate this, we compared LINSIGHT scores between simple and complex enhancer architectures. Briefly, LINSIGHT estimates the probability of negative selection on noncoding sites in the human lineage at a base-pair level using both functional genomics annotations and conservation metrics; higher scores indicate stronger negative selection (Huang et al., 2017). In general, complex enhancers have higher LINSIGHT scores than simple enhancers, suggesting slightly stronger negative selection in complex enhancers (0.16 versus 0.14 mean LINSIGHT score; *p* < 2.2e-308). Given that simple and complex enhancer sequences have different age distributions (Figure 3C, Supplemental Figure 3.3), we stratified enhancers by age to evaluate whether simple enhancers consistently had lower scores than complex enhancers. This revealed that simple enhancers generally have similar LINSIGHT scores as age-matched complex enhancers. Overall, these findings are consistent in our analysis of Roadmap enhancers (Supplemental Figure 3.3). Similarly, analysis of simple and complex enhancer PhastCons element overlap supports that complex enhancers are overall more conserved than simple enhancers and that the majority of both simple and complex enhancers are highly conserved at older ages (Supplemental Figure 3.1). Together, these results suggest that complex enhancers on average experience stronger purifying selection than simple enhancers, though these differences are likely due to the differences in sequence age distributions. Age-matched simple and complex enhancers have similar selection estimates, indicating both architectures experience similar selection pressures if they survive the evolutionary process.

### Genetic variants in simple enhancers are more likely to be associated with human traits and disease than variants in complex enhancers

The majority of genetic variants associated with human complex traits and disease are located in functional, non-coding regulatory regions (Corradin & Scacheri, 2014; Maurano et al., 2012). Based on the differences in pleiotropy and constraint observed between architectures, we hypothesized that enhancer architecture could provide evolutionary context for interpreting the effects of enhancer variants on traits. To test this, we evaluated enrichment of 55,480 significant (*p* < 5e-8, linkage disequilibrium expanded at r^2^=1) GWAS Catalog single-nucleotide variants from 2,619 genome-wide association studies (Buniello et al., 2019) in simple and complex architectures compared to length- and architecture-matched background regions. We observe a strong enrichment in simple enhancers and a depletion of variants in complex enhancers compared with expected levels (Figure 4A, 6.8-fold-change for simple versus 0.6-fold-change complex architecture; *p* < 2.2e-308, Welch’s Test). This difference also held when stratifying enhancers by age (Supplemental Figure 4.1). We also tested for GWAS variant enrichment in Roadmap ChIP-peak-defined enhancers by trimming them to the center 310 bp (Supplemental Figure 2.10, Methods). We found enrichment for GWAS variants overall, but no consistent tissue-by-tissue difference in the enrichment among simple and complex enhancers (median 1.12-fold-change in simple versus 1.13-fold-change in complex; *p* = 0.28, Mann Whitney U; Supplemental Figure 4.2). However, we did not attempt to match the GWAS considered to the different tissue contexts; thus, more work is needed to evaluate variation in simple and complex enhancer enrichment across tissues.

**Figure 4.**
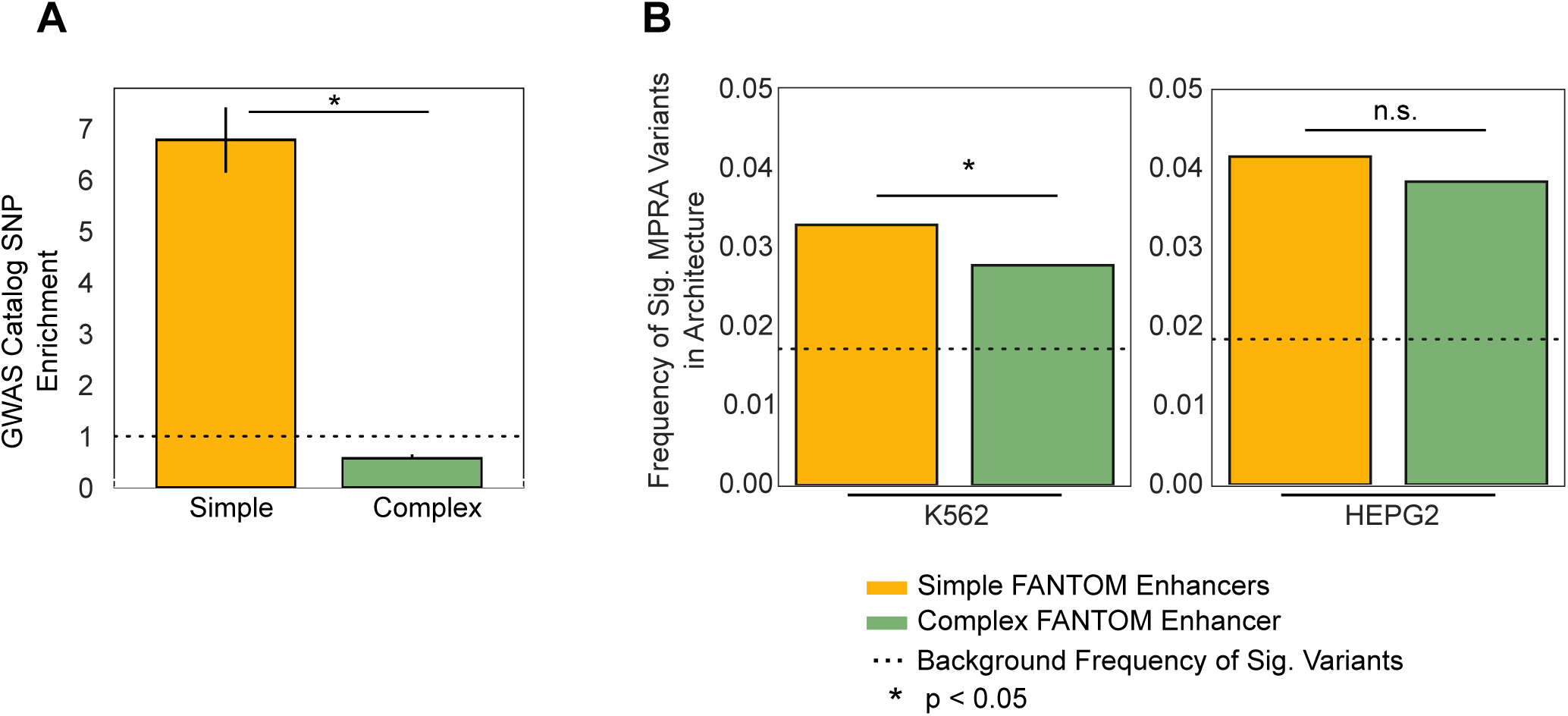
Simple enhancers are enriched for GWAS variants and variants with significant regulatory activity in massively parallel reporter assays. (A) Simple and complex enhancers are both significantly enriched for GWAS catalog variant overlap (6.8-fold enrichment in simple enhancers versus random regions, FDR 5% *p* = 0.01, and 0.6-fold enrichment in complex enhancers versus random regions, FDR 5% *p* = 0.01). Simple enhancers are more enriched for GWAS variants than complex enhancers (*p* < 2.2e-308, Welch’s Test). Enrichment was estimated based on comparison to the distribution of overlaps observed in random permuted enhancer sets; error bars represent 95% confidence intervals based on 10,000 bootstraps. (B) Genetic variants in simple enhancers are enriched for significant changes in regulatory activity compared to variants in complex enhancers in massively parallel reporter assays (MPRAs) in K562 cell models (odds ratio (OR) = 1.18; *p* = 0.04, Fisher’s Exact Test) and in HepG2 cell models (OR = 1.09; *p* = 0.26). The fraction of significant alleles was calculated from all tested variants overlapping simple and complex enhancer architectures and is plotted on the y-axis. In K562 cell models, 3.3% of variants in simple enhancers and 2.6% of variants in complex enhancers exhibit significant changes in MPRA activity compared to 1.7% of background variants (simple OR = 1.92, *p* = 2.13e-35, and complex OR = 1.62, *p* = 1.69e-10). In HepG2 cell models, 4.1% of variants in simple enhancers and 3.8% of variants in complex enhancers produce significant changes in MPRA activity compared to 1.8% of background variants (simple OR = 2.30, *p* = 1.57e-66 and complex OR =2.12, *p* = 3.75e-29). The dashed horizontal lines represent the background fraction of all variants tested with significant activity per cell line. Significant activity was called at a 5% FDR threshold.

To explore the patterns of clinically relevant variants in different enhancer architectures, we evaluated ClinVar disease-associated variant enrichment in simple and complex enhancers (Landrum et al., 2018). While GWAS associations reflect variant effects on common, complex diseases, pathogenic variants in ClinVar are often involved in rare Mendelian disorders. Simple enhancer variants were enriched for “pathogenic” annotations in both eRNA and ChIP-peak datasets, while complex enhancers were enriched for “benign” annotations in both datasets, yet only complex ChIP-peak enhancers were significantly enriched (median 29.0% complex versus 27.8% simple enhancers overlap “benign” ClinVar variants; *p* = 1.1e-4, Mann Whitney U, Supplemental Figure 4.4). Together, these results confirm enrichment for trait and rare disease variants in both complex and simple enhancer architectures compared to regions without enhancer activity; however, known trait-associated variation occurs more frequently in simple enhancer architectures, while benign variation occurs more frequently in complex architectures.

To complement these findings, we evaluated the enrichment of known expression quantitative trait loci (eQTL). Simple and complex enhancers were similarly enriched for GTEx eQTL across 46 tissues (GTEx Consortium, 2017) at ∼1.1x fold-change (Supplemental Figure 4.3; median 1.09 and 1.11 simple and complex enhancer fold change; *p* =0.38, Mann Whitney U). This indicates that both architecture types are similarly likely to contain variants associated with gene expression variation across individuals.

### Genetic variants in simple enhancers are enriched for changes in biochemical regulatory activity compared to variants in complex enhancers

Given the differences in genetic variant associations with organism-level traits between simple versus complex enhancers, we hypothesized that there would be architecture-related differences in the effects of variants on gene regulatory activity at the molecular level. We tested for enrichment of variants that significantly affect biochemical regulatory activity among trimmed simple and complex architectures. We considered >30,000 common human variants shown to affect regulatory activity in recent massively parallel reporter assays (MPRA) performed in K562 and HepG2 cells (van Arensbergen et al., 2019). For both cell lines, variants in annotated enhancers were significantly more likely to have regulatory effects than all background variants tested in the assay (Figure 4B, simple OR = 1.92, *p* = 2.13e-35 Fisher’s exact test in K562 and OR = 2.30, *p* = 1.57e-66 in HepG2; complex OR = 1.62, *p* = 1.69e-10 in K562 and OR =2.12, *p* = 3.75e-29). Simple architectures are more enriched than complex architectures for variants that significantly affect regulatory activity in both K562 (OR = 1.18, *p* = 0.04) and in HepG2 cells, although the enrichment is smaller in HepG2 cells (OR = 1.09, *p*= 0.26). Trimmed Roadmap K562- and HepG2-defined enhancers are also enriched for significant MPRA activity, and simple enhancers are more enriched, but the enrichments are modest (Supplemental Figure 4.5, OR = 1.26, *p* = 0.13 in HepG2 assays, and OR = 1.25 and *p* = 0.18 in K562 assays). We repeated this analysis using only granulocyte and liver FANTOM enhancers to match the cellular contexts tested and found even stronger enrichment among simple enhancers in these datasets (liver OR = 1.8; *p* = 0.08 and granulocyte OR = 1.32; *p* = 0.13, Fisher’s exact test) (Supplemental Figure 4.5). These findings indicate that common human variants in simple enhancers are more likely to significantly affect enhancer biochemical regulatory activity.

### Simple enhancers overlap transposable element derived sequences more often than complex enhancers

TE-derived sequences (TEDS) have enhancer activity across many cellular contexts (Chuong et al., 2017; Marnetto et al., 2018; Simonti et al., 2017; Su et al., 2014; Sundaram et al., 2014; Trizzino et al., 2017). A previous study identified that TE insertions occur nearby sequence age breaks (Marnetto et al., 2018). We hypothesized that TEDS might have different influences on simple and complex enhancer architectures. To explore this, we tested TEDS enrichment in enhancer architectures against the genomic background. To control for length differences, we evaluated both 310 bp and 1 kb trimmed/expanded enhancers. Both length-control strategies yielded similar results, and we present the 310 bp results below. We intersected the 310 bp enhancers with genome-wide maps of TEDS (Methods). We find that 48% of simple enhancers and 42% of complex enhancers contain TEDS. As expected from previous reports (Emera et al., 2016; Simonti et al., 2017), both simple and complex enhancers are depleted of TEDS compared to architecture-matched genomic backgrounds, and complex enhancers are substantially more depleted (Figure 5A, OR = 0.50 versus 0.25; *p* < 2.2e-308, Fisher’s exact test). The majority of enhancer sequences younger than the eutherian ancestor contain TEDS (Figure 5C). In complex enhancers, TEDS are often younger than the therian MRCA, while for simple enhancers, most are younger than the eutherian ancestor (Supplemental Figure 5.3). This is consistent with previous observations that the majority of young human/primate cis-regulatory elements contain TEDS (Simonti et al., 2017; Trizzino et al., 2017).

**Figure 5.**
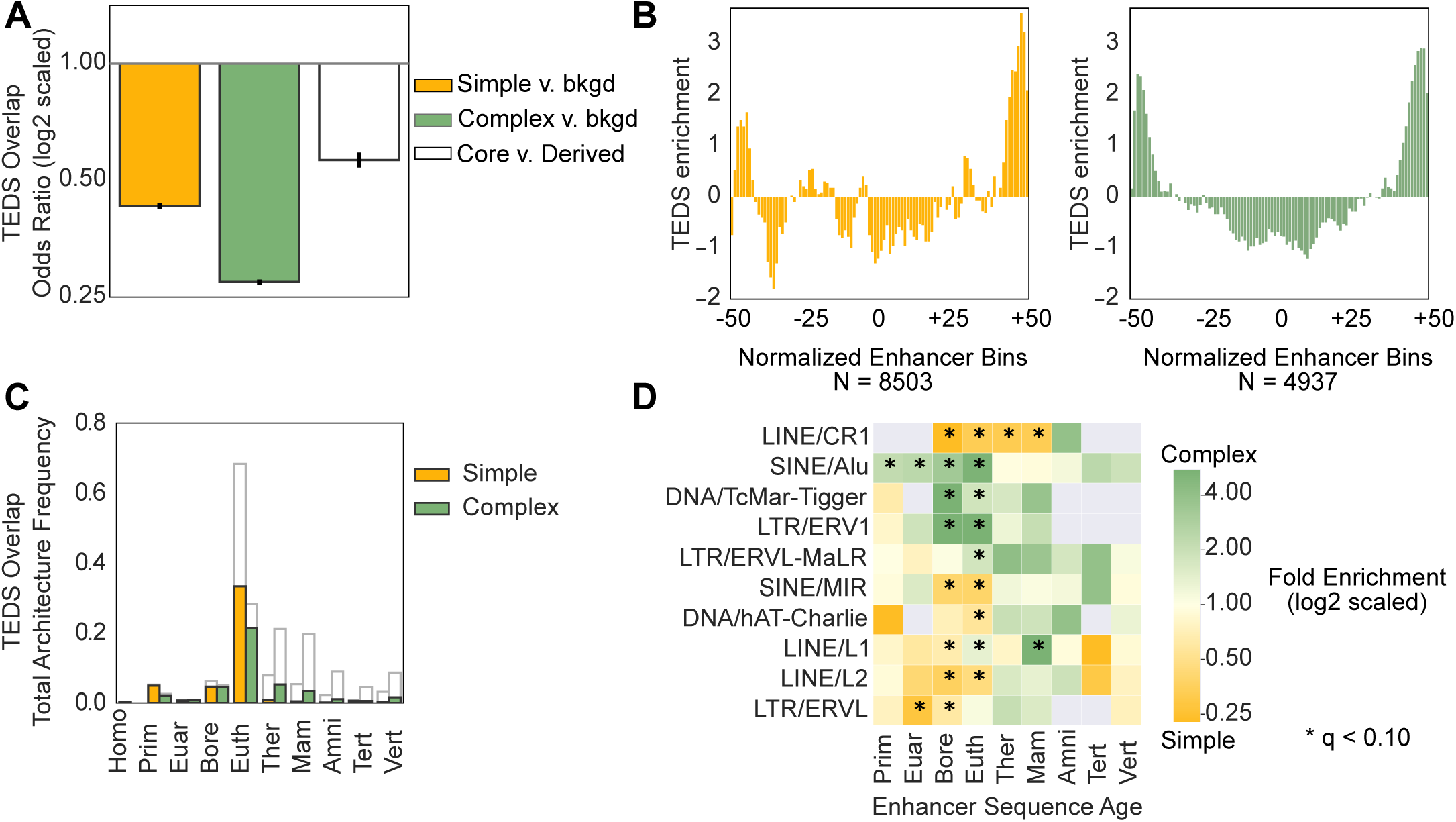
Simple and complex enhancers are enriched for sequences derived from different transposable element families across ages. (A) Simple and complex enhancers are significantly depleted of transposable element derived sequences (TEDS) compared to 100 architecture-matched random controls (∼2-fold depleted for complex and ∼4-fold depleted for simple; *p* <2.2e-308, Fisher’s Exact Test). Within complex enhancers, older core sequences are ∼2 fold-depleted of TEDS compared with the younger, derived sequences (OR = 0.56; *p* = 9.72e-89). (B) TEDS are enriched in the outer 50% of complex enhancer sequences. TE enrichment is the z-score of TE overlap counts in each normalized enhancer bin across complex enhancers (green, median z-score = 0.17 versus -0.73, outer v inner 50% bins; *p* = 6.38e-18, Mann Whitney U) and simple enhancers (yellow, median z-score = -0.43 versus -0.43, outer v inner 50% bins; *p* = 0.47). (C) Fraction of all simple and complex enhancers overlapping TEDS stratified by age. Grey outlines represent overall fraction of simple or complex enhancers of each age. (D) Simple and complex enhancers are significantly enriched for sequences derived from different TE families across ages. TEDS enrichment in enhancer architectures was calculated among TEDS-overlapping enhancers of each age using Fisher’s exact test. Positive values (green) represent TEDS enrichment in complex enhancers, while negative values represent enrichment in simple enhancers (yellow, FDR *p* < 0.10).

### Transposable element sequences can both nucleate and remodel enhancers

Sequences with some regulatory potential have been hypothesized to nucleate enhancer activity, which can then be expanded and remodeled by the addition of younger sequences (Emera et al., 2016). To explore the role of TEDS in this process, we tested for TEDS enrichment in complex enhancer core sequences versus younger derived sequences. Overall, complex enhancer cores are depleted of TEDS compared with derived sequences (Figure 5A, Supplemental Figure 5.1, OR = 0.56; *p* = 9.7e-89). We also found strong enrichment for TEDS at the edges of complex enhancers and depletion at their centers (Figure 5B, green, median z-score = 0.17 versus -0.73, outer v inner 50% bins; *p* = 6.4e-18, Mann Whitney U). These results are consistent with our finding that younger sequences flank older core sequences, and suggests that TEDS often contribute younger sequences to complex enhancer architectures. However, this general trend is largely driven by old complex enhancers; younger complex enhancers (younger than the therian MRCA) are enriched for TEDS in their cores (Supplemental Figure 5.2). By comparison, TEDS are also enriched at the edges of simple enhancers, though the central regions of simple enhancers do not show strong TEDS depletion (Figure 5B right panel Supplemental Figure 5.1). These results support a model where TEDS can both nucleate and remodel enhancer sequences.

### Enhancer architectures are enriched for different TE families

As discussed above, TE insertions can disrupt functional elements and lead to genome instability. Thus, the probability of TE insertions gaining gene regulatory activity likely depends on their genomic sequence context. We hypothesized that enhancers with different architectures and origins would be enriched for TEDS from specific TE families. Several TE families show biases for simple or complex enhancer architectures at different evolutionary ages (Figure 5D). Complex enhancers are consistently enriched across ages for SINE/Alu, DNA/TcMar-Tigger, LTR/ERVL-MaLR, and LTR/ERV1 elements. SINE/Alu elements are abundant in the primate lineage (Batzer & Deininger, 2002), but are frequently observed in complex enhancers older than the primate ancestor. Integrating young SINE/Alu TEDS with these older sequences may have altered ancient regulatory activity or created new regulatory activity. Simple enhancers are consistently enriched across ages for LINE/CR1, SINE/MIR, DNA/hAT-Charlie, LINE2/L2, and LTR/ERVL elements (Figure 5D). LINE/L1 elements are significantly enriched in both older complex and younger simple enhancers, suggesting L1s contribute sequence to both architectures. Together, these data suggest differences in the contribution of TEs to enhancer sequences of different origins and sequence architectures.

## DISCUSSION

Here, we evaluate the genomic, evolutionary, and functional features associated with human enhancers of different sequence age architectures. As seen in previous work, the majority of sequences with human enhancer activity originate from the placental MRCA or older ancestors. We expand beyond considering the age of an enhancer sequence by defining its sequence age architecture. Human enhancers have many distinct age architectures, and we demonstrate that simple architectures are favored over complex architectures. Functionally, simple and complex architectures show differences in tissue-specific and cross-species activity profiles. However, both architectures experience similar selective constraints as they age. Simple architectures are more enriched for variants associated with complex traits in GWAS studies, rare pathogenic variants in ClinVar, and variants that significantly alter biochemical activity. Sequences derived from TEs are depleted among all enhancers, but they are more depleted in complex architectures than simple. Nonetheless, these TEDS provided genomic material for many younger enhancers of both architectures and modified older sequences into complex architectures with enhancer activity. Distinct TE families are enriched in different architectures. Thus, TEDS have made important contributions to the evolution of both simple and complex sequence age architectures. Finally, the consistency of these observations across enhancer sequences identified from eRNA signatures and histone modification patterns supports their generality.

Our work expands our current understanding of enhancer sequence evolution in several dimensions. We show that features of the two-step proto-enhancer life-cycle model proposed by Emera et al. are present in enhancers across diverse tissues. However, the depletion for complex architectures among enhancer sequences suggests that this model does not represent the most common history for human enhancer sequences. Several lines of evidence suggest that human simple enhancers are not simply a snapshot of proto-enhancers in the first step of the enhancer life cycle: (1) Simple enhancer sequences are often as old as complex enhancers, implying that these ancient sequences did not require major remodeling to obtain gene regulatory functions. (2) Simple and complex enhancers of similar ages are under similar levels of purifying selection. (3) Simple enhancers are enriched for tissue-specific functions. (4) Simple enhancers are enriched for GWAS variants, pathogenic ClinVar variants, and variants impacting biochemical activity, implying that simple enhancer variation contributes to human trait variation and changes in molecular function. Together, these results suggest that simple enhancers play important roles in human gene regulatory biology. However, we speculate that simple enhancer sequences may be less evolutionarily stable, as fewer older simple enhancers are observed. In contrast, complex enhancers may be more functionally robust to mutations given their older ages, increased cross-species activity, and trait variant patterns. Further, complex enhancers contain younger derived sequences that are under less selective pressure and expand enhancer length. These younger derived sequences are enriched for eQTL and higher minor allele frequencies across diverse human populations (data not shown), which agrees with findings that functional stability of TF binding motif mutations is influenced by flanking sequences (Li et al., 2019). We speculate that younger derived sequences could protect complex enhancers from inactivating mutations. Future biochemical work can address whether architectural features of complex enhancers may make them more robust to mutations.

To integrate our findings and provide a framework for future work, we propose a model for the evolution of enhancer sequences architecture and activity (Figure 6). In our model, inspired by Markov models, sequences occupy either simple or complex architecture states and either active or inactive states. Genomic events (e.g., substitutions and rearrangements) drive transitions between these states over time. Based on our results, we propose that certain paths through the model are common in the enhancer life cycle. Most sequences that ultimately obtain enhancer activity likely begin as inactive or weakly active sequence segments (Figure 6, left). Small-scale genomic events, like point mutations, can strengthen regulatory activity and create simple enhancers (Figure 6, top right). Examples include human accelerated regions, such as HACNS1/HAR2, where human-specific substitutions have created human-specific enhancer activity in limb bud formation (Cotney et al., 2013; Prabhakar et al., 2008). TE insertions also give rise to simple enhancers by integrating sequence with regulatory potential into genomes (Chuong et al., 2017); for example, the mouse-specific RLTR13 endogenous retrovirus sequence is sufficient to drive gene expression in rat placental cells (Chuong et al., 2013).

**Figure 6.**
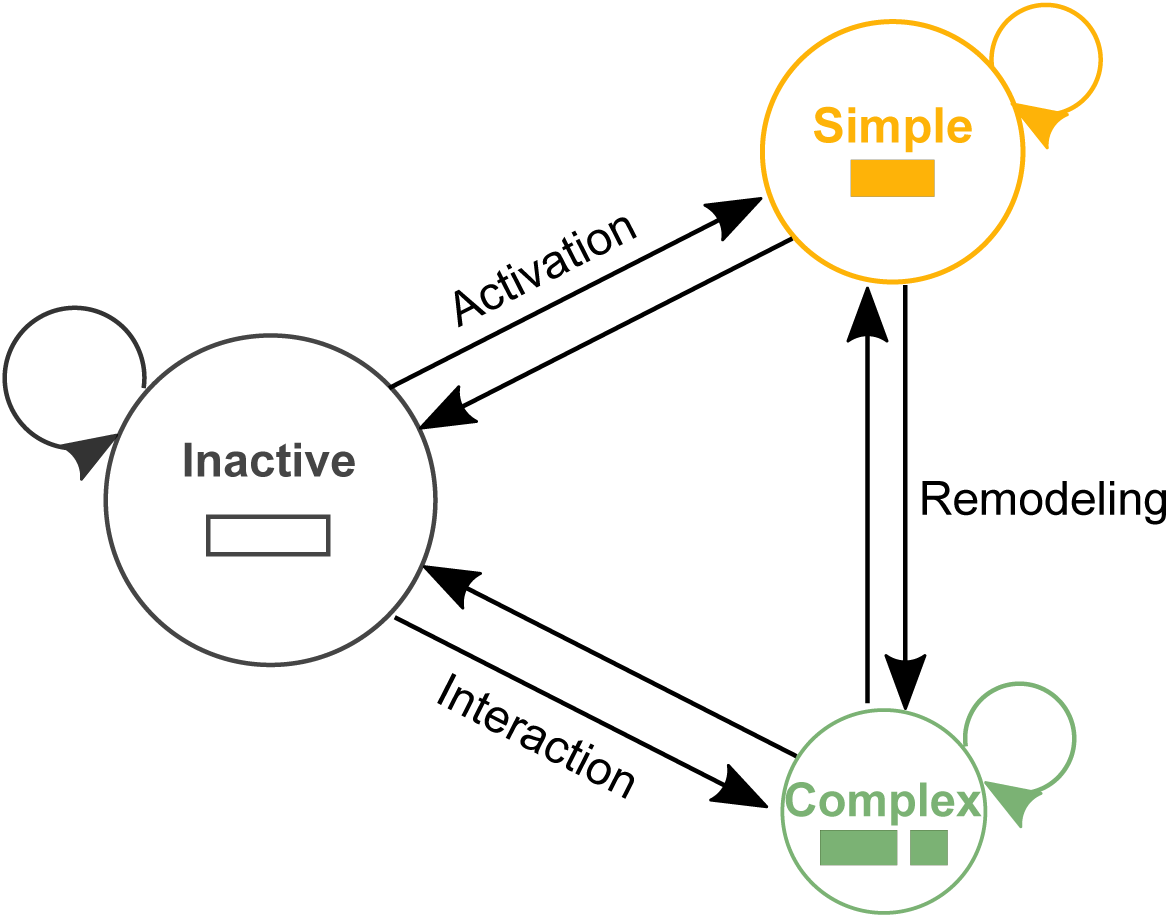
Model of enhancer sequence architecture and activity. Sequences with the potential to be enhancers can occupy active or inactive states and have either simple or complex sequence age architectures. Sequences transition between these states as a result of large- and small-scale genomic variants. Inactive sequences can become simple enhancers through small-scale genomic changes, such as substitutions that increase activity or nearby chromatin changes that increase accessibility. Complex enhancers consist of sequences of different origins brought together by genomic rearrangements. In some cases, the integration of these sequences and subsequent substitutions produce activity, and in others already active simple enhancers are remodeled into active complex enhancers with different activity patterns. Sequences regularly transition between these states over evolutionary time. Sequences with human enhancer activity are enriched for simple evolutionary architectures.

Complex enhancers can emerge from multiple different evolutionary paths. For example, large-scale (greater than a few nucleotides) genomic insertions or rearrangements combined with small-scale substitutions may remodel active simple enhancers into complex enhancers with stronger or different activity patterns (Figure 6 right). Work in *Drosophila* has demonstrated that small-scale substitutions in complex cross-vein and wing spot enhancers “co-opt” ancestral enhancer activity to develop lineage-specific wing pigmentation patterns (Koshikawa et al., 2015; Prud’homme et al., 2006). Isolated derived segments in these complex enhancers were not sufficient to drive enhancer activity during development, but may function to support lineage-specific enhancer activity in other ways, such as facilitating cooperative or co-activator binding (Long et al., 2016). Complex enhancers can also be created when genomic rearrangements place weakly active sequences of different origins adjacent to each other in such a way that these sequences interact and/or accumulate additional substitutions to create a new active complex enhancer (Figure 6 bottom right). TE insertions can facilitate such interactive effects. For example, the interaction of a LINE/L2 insertion and flanking sequence formed a new enhancer that was both necessary and sufficient for driving increased, lineage-specific GDF6 expression and evolutionary changes in armor-plate size in freshwater stickleback (Indjeian et al., 2016). TE insertion near older active regulatory sequences can protect TEDS from inactivation by the host genome, creating substrates for complex enhancers to form (Elbarbary et al., 2016; Levin & Moran, 2011; Varshney et al., 2015). Without further experimental dissection, it is challenging to trace the history of activity, especially for complex enhancer sequences. For example, human-specific substitutions in human accelerated region 2xHAR.164 have remodeled its enhancer activity from the spinal cord to other regions of the developing brain (Capra et al., 2013a). Analysis of the 2xHAR.164 locus reveals that the underlying complex enhancer sequence is composed of an amniote core and a eutherian derived segment. If the amniote core functioned as an enhancer without the derived eutherian sequence, a remodeling event might have occurred in the past. Alternatively, if amniote and eutherian sequences in isolation do not have enhancer activity, then an interaction might have occurred.

We emphasize that most enhancer sequences do not reach a final stable state; sequences continue to change. Thus, we constructed our model (Figure 6) to emphasize that sequences regularly transition between these states over evolutionary time. Large comparative regulatory genomics datasets across species and tissues are needed to estimate these transition probabilities. We hope that future work will enable us to measure rates of simple and complex enhancer emergence, decay, and turnover across other species.

Several limitations must be considered when interpreting our results. First, sequence age estimates are influenced by the accuracy of sequence alignment methods, genome quality, and different rates of sequence divergence across the genome over evolutionary time (Capra et al., 2013b; Cooper & Brown, 2008; Margulies & Birney, 2008). Assembling and aligning repetitive elements is particularly challenging and may limit TEDS detection (Ewing, 2015). Thus, our estimates should be viewed as lower bounds on the actual sequence age. Second, our analyses are limited by the availability of enhancer datasets. Our tissue pleiotropy estimates reflect the adult enhancer landscape and do not sufficiently survey developmental or stimulus-dependent enhancer landscapes; these should be evaluated separately in future work. Further, enhancer resolution in tissue-level datasets is complicated by underlying cell heterogeneity. Our cross-species activity analysis is limited to liver enhancers from nine species. Third, we are limited in our knowledge of human-trait and disease-associated variants. GWAS-variant enrichment reflects tag-SNP and LD-linked loci associated with measurable common human traits; we do not directly demonstrate that they contribute to disease pathology or affect enhancer activity. ClinVar-variant enrichment is limited by the small number of known pathogenic non-coding variants. Some analyses are underpowered, and as a result, we did not find statistically significant associations between simple architectures and pathogenic variants in either dataset. Finally, we do not analyze sequence-level features that distinguish simple and complex architectures, and thus are limited in our descriptions of such features (e.g., distinct TF binding sites) that may have facilitated the evolution of enhancer architectures. We envision that a thorough analysis of binding site motifs and sequence ages in these architectures will elucidate the determinants of evolutionary trajectories in enhancer sequence architectures.

In conclusion, we related sequence age architectures to human enhancer function and genetic variation. Evaluating architecture revealed different evolutionary origins and dynamic evolutionary trajectories among enhancer sequences. We present a model of enhancer sequence evolution, which encompasses these multiple evolutionary trajectories. Our work provides a foundation for future studies that dissect the relationships between enhancer sequence architecture, evolution, and the consequences on function and non-coding variation in the human genome.

## METHODS

### Syntenic block aging strategy

The genome-wide hg19 46-way vertebrate multiz multiple species alignment was downloaded from the UCSC genome browser. Each syntenic block was assigned an age based on the most recent common ancestor (MRCA) of the species present in the alignment block in the UCSC all species tree model (Figure 1A). For most analyses, we focus on the MRCA-based age, but when a continuous estimate is needed we use evolutionary distances (in substitutions per site) from humans to the MRCA node in the fixed 46-way neutral species tree. Trees were analyzed using the python package, dendropy (Sukumaran & Holder, 2010). Estimates of the divergence times of species pairs in millions of years ago (MYA) were downloaded from TimeTree (Hedges et al., 2015). Sequence age provides a lower-bound on the evolutionary age of the sequence block. Sequence ages could be estimated for 93% of the base pairs (bp) in the human genome.

### Enhancer aging and architecture assignment

We considered enhancers called from enhancer RNAs (eRNAs) identified across 112 tissue and cell lines by high-resolution cap analysis of gene expression sequencing (CAGE-seq) carried out by the FANTOM5 consortium (Andersson et al., 2014). This yielded a single set of 30,474 autosomal enhancer coordinates. We assigned enhancer ages by intersecting their genomic coordinates with aged syntenic blocks using Btools v2.27.1 (Quinlan & Hall, 2010). Syntenic blocks that overlapped at least 6 bp of an enhancer sequence were considered when assigning the enhancer’s age and architecture. We considered enhancers mapping to one syntenic block or several syntenic blocks of the same age as “simple” enhancer architectures, while enhancers overlapping adjacent syntenic blocks of different ages have “complex” enhancer architectures. Given complex enhancers are composed of multiple sequence ages, we assigned complex enhancer age according to the oldest age. Sequences without an assigned age were excluded from this analysis.

We also considered enhancers identified by the Roadmap Epigenomics Mapping Consortium (Roadmap Epigenomics Consortium et al., 2015) across 98 cellular contexts. Roadmap defined enhancers from histone modification chromatin immunoprecipitation (ChIP-seq) peaks by subtracting H3K4me3+ peaks from H3K27ac+ peaks to exclude active promoters. This resulted in 20,172,949 predicted autosomal enhancers. Enhancers <10 kb in length were considered. Roadmap enhancers were assigned ages and architectures as described above for the FANTOM enhancers.

From the human syntenic blocks that could be assigned ages, the plurality (44%) are derived from the placental MRCA, while 40% are younger than the placental MRCA, and 16% are older (Supplemental Figure 1.1A). This result was consistent with syntenic age estimates using hg38 and 100-way species alignments (Marnetto et al., 2018). Younger syntenic blocks are generally longer than older syntenic blocks (median 128 bp for primate-specific blocks versus 42–66 bp for older syntenic blocks) (Supplemental Figure 1.1B).

Enhancers are often defined by multiple different strategies that have low levels of overlap and no single strategy captures the breadth of active enhancers (Benton et al., 2019). Thus, the general concordance in enhancer architecture patterns among enhancers defined by eRNA and histone-based ChIP-seq experiments is reassuring.

### Trimming and expansion of enhancer lengths

We trimmed or expanded Roadmap enhancers to 310 bp to equalize enhancer lengths between ChIP-seq and eRNA sets. Other methods, such as DNase sensitivity, more precisely identify sequences with enhancer activity than the entire sequence length underlying a ChIP-seq peak (Andersson & Sandelin, 2020). However, trimming ChIP peak sequences has limitations. First, it assumes peak centers represent the most active segment of the enhancer sequence. Second, we exclude flanking sequences that may be important for opening chromatin or recruiting transcriptional machinery. Third, it may bias analysis of complex enhancers towards older sequences, as older sequence ages tend to occur at enhancer centers. Finally, multiple active enhancer sub-regions might be dispersed throughout a peak or constitute super-enhancers. To explore the sensitivity of our results on Roadmap enhancers to this choice, we also analyzed the entire (median ∼2.4 kb) ChIP-seq peaks. Single syntenic blocks of this length are rare in the human genome, thus simple enhancers are rare when considering the full peak. Nonetheless, we still observed significantly fewer segments of different ages among the full Roadmap enhancers (Supplemental Figure 2.5A, Supplemental Figure 2.10A).

### Human syntenic block PhastCons conservation

PhastCons vertebrate hg19 conserved elements were downloaded from the UCSC genome browser (Siepel, 2005). PhastCons elements were assigned ages using the same MRCA-based strategy described for enhancers. As expected, sequence age is correlated with sequence conservation (r^2^ = 0.82, *p* = 0.009), since sequence homology is the basis for estimating both sequence age and sequence conservation. However, these metrics capture complementary information about regions of interest. Sequence conservation summarizes the evidence that purifying selection has acted on the region, and conserved sequences have high similarity across species. Sequence age estimates a lower bound on the evolutionary origin of a sequence and can be assigned both to conserved sequences and neutrally evolving sequences with lower sequence identity among species. For example, only 35% of the oldest syntenic blocks have significant evidence of evolutionarily conservation (vertebrate PhastCons overlap, Supplemental Figure 1.1 C). In other words, not all old sequences have evidence of significant conservation. Thus, even though neutrally evolving sequences become more difficult to accurately age with time (such that age reflects a lower bound estimate of sequence origin), sequence age provides complementary information about sequences shared among vertebrates.

### Background random genome regions and architectures

For FANTOM enhancers, 100 random shuffles of the genomic regions in each dataset of interest (e.g., cellular context) were performed using BEDTools. For Roadmap enhancers, each of the 98 tissue datasets was shuffled 10 times, resulting in 980 shuffled datasets total. The shuffled sets were matched on chromosome number and length, and they excluded ENCODE blacklist regions and genomic gaps as defined by the hg19 UCSC gaps track (Amemiya et al., 2019). Random genomic regions were then assigned ages and architectures with the same strategy used for enhancers described above. We calculated enrichments by comparing the observed enhancer age and architecture distribution with the expectation from the appropriate sets of shuffled regions.

### Enhancer tissue pleiotropy

To account for the effects of enhancer length in quantification of enhancer activity across biological contexts. FANTOM enhancers were trimmed around their midpoints to the mean length of all enhancers in the dataset (310 bp), and Roadmap enhancers were similarly trimmed to 310 bp to control for differences in length between eRNA and ChIP-seq enhancers. Trimmed enhancer datasets were intersected with 112 FANTOM eRNA tissue facets and cell line datasets or with 97 Roadmap ChIP-seq datasets using BEDTools multi-intersect command. We considered an enhancer pleiotropic when at least 50% of the enhancer length overlapped enhancers in other contexts.

### Cross-species enhancer activity

Human liver enhancers from a cross-species analysis of vertebrate livers (Villar et al., 2015) were assigned ages and architectures. To account for length differences, complex enhancer length was randomly matched to the simple enhancer lengths (N = 1308 and N = 1520 matched-length complex and simple enhancers). The number of species in which the sequence is present and has enhancer activity was determined from pairwise sequence alignments of enhancer maps from nine placental mammals.

### Enhancer sequence constraint

LINSIGHT scores were downloaded from http://compgen.cshl.edu/~yihuang/LINSIGHT/. LINSIGHT provides per base pair estimates of negative selection (Huang et al., 2017). Enhancers were intersected with LINSIGHT base pair estimates. 46-way hg19 vertebrate PhastCons elements were downloaded from the UCSC genome browser. Enhancers overlapping any PhastCons element by at least 6 bp were considered conserved.

### GWAS catalog enrichment

Enrichment for overlap with 55,480 GWAS Catalog variants (p<5e-8) from 2601 traits (last downloaded September 24rd, 2019) (Buniello et al., 2019) were linkage disequilibrium expanded (r^2^ =1.0) using European 1000 Genome phase reference panels (The 1000 Genomes Project Consortium, 2015). Enrichment was tested by comparing the observed overlap for a set of regions of interest with overlaps observed across 100 shuffled sets matched on length, sequence age architecture, and chromosome. Median fold-change was calculated based on the GWAS Catalog variants overlapping enhancer architectures compared with these random genomic sets. Confidence intervals (CI = 95%) were generated by bootstrapping the 1000 random genomic fold-change values 10,000 times. P-values were corrected for multiple hypothesis testing by controlling the false discovery rate (FDR) at 5% using the Benjamini-Hochberg procedure.

### ClinVar variant enrichment

ClinVar variants in VCF format were downloaded from ftp://ftp.ncbi.nlm.nih.gov/pub/clinvar/ (last downloaded 2019-12-02). Trimmed FANTOM and Roadmap enhancers were intersected with ClinVar variants. FANTOM enhancers overlapped 21 annotated variants total (n=9 simple, n=12 complex). Among 98 Roadmap tissue enhancer sets, enhancers overlapped a median of 6690 annotated ClinVar variants (range = 1590 to 22797 variants total). ClinVar variants were considered pathogenic if annotated with the term “pathogenic” and excluded if annotated with the term “conflicting”. Similar inclusion and exclusion criteria were used for “benign” and “protective”. The fraction of annotated variants per architecture was estimated as the number of “pathogenic”, “benign”, or “protective” annotations versus all ClinVar variants overlapping that architecture.

### eQTL enrichment

Enrichment for GTEx v6 eQTL from 46 tissues (last downloaded July 23rd, 2019) (GTEx Consortium, 2017) in enhancers with simple and complex architectures was tested against a null distribution determined by shuffling observed enhancers using the same strategy as described for GWAS variant enrichment.

### Massively parallel reporter assay data

Results from MPRAs were downloaded from van Arensbergen 2019 (van Arensbergen et al., 2019). Significant changes in MPRA activity and p-values were calculated by the authors using a Wilcoxon rank-sum test with an FDR = 5% separately identified in K562 and HepG2 cell lines. Trimmed enhancers were intersected with alleles tested in MPRA. Fisher’s exact test was used to estimate the odds an allele with significant changes in MPRA activity occurred in a specific architecture compared with the background set of variants that do not overlap enhancers. Significant allele overlap was also compared between simple and complex enhancer architectures to estimate an odds ratio of enrichment.

### Transposable element derived sequence enrichment

Transposable element derived sequences identified by RepeatMasker were downloaded from the UCSC genome browser and liftedOver to hg19 from hg38 (last downloaded 4-14-2018). Trimmed enhancers (310 bp) were intersected with TEDS coordinates. TEDS overlapping enhancers >= 20 bp were evaluated further for enrichment in FANTOM enhancers of different ages. Enrichment was estimated as the number of TEDS in enhancer architectures compared with random-shuffled regions matched on both length and architecture using Fisher’s exact test. We compared enrichment between core and derived segments of complex enhancers by using Fisher’s exact test on TEDS overlap counts in core and derived syntenic blocks. To estimate TEDS family enrichment in enhancers with different sequence age architectures, we compared the number of simple/complex enhancers overlapping a TEDS family with the number of simple/complex architectures overlapping any other TEDS family of that age. Enrichment significance was evaluated using Fisher’s exact test and FDR controlled at 10%.

