## supplemental_figures for "Modeling the evolutionary architectures of human enhancer sequences reveals distinct origins, functions, and associations with human-trait variation"

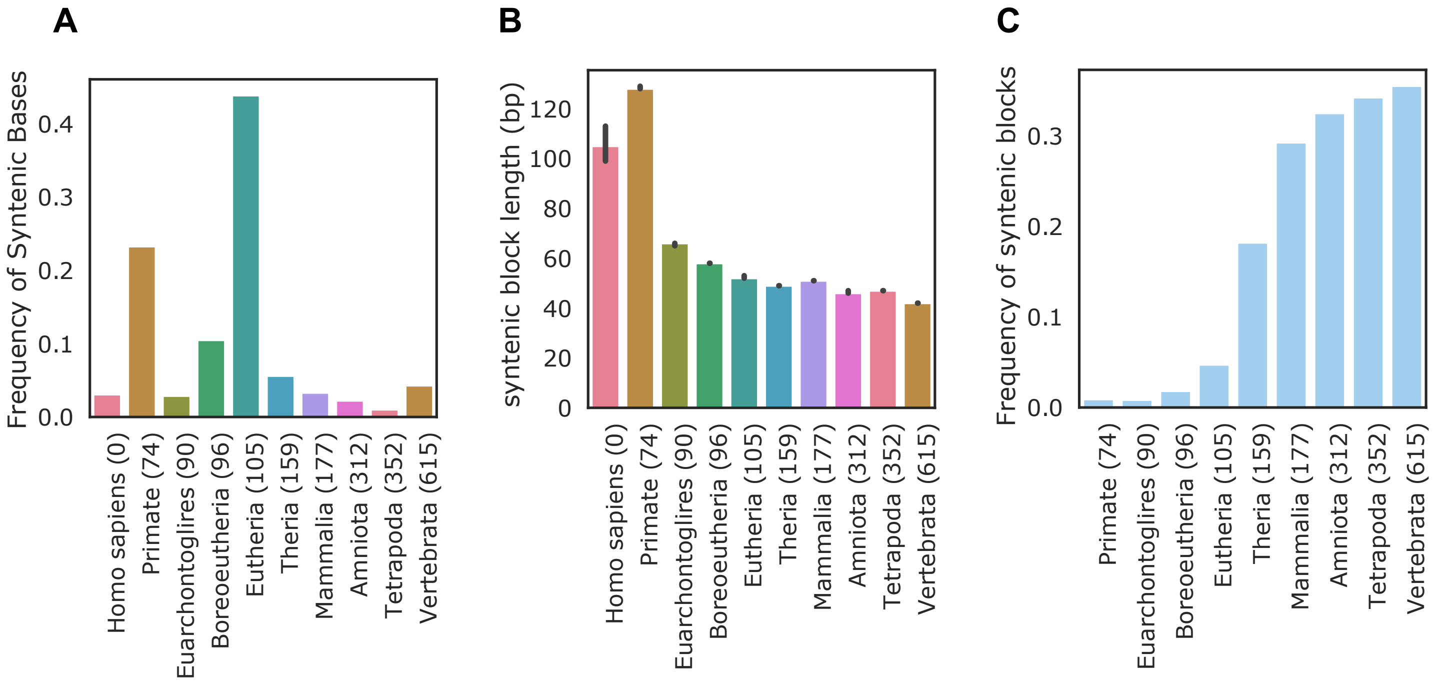


**Supplemental Figure 1.1. Human genome age distribution.** (A) Most human syntenic blocks are derived from the placental MRCA. Sequence age for each base pair from hg19 UCSC 46-way MultiZ sequence alignments. Sequence age are assigned to each syntenic block based on the oldest most recent common ancestor (MRCA) of extant species alignable with humans. (B) Younger syntenic blocks are longer than older syntenic blocks. Median syntenic block length per age. (C) A minority of syntenic blocks per age overlap PhastCons elements. Percent of syntenic blocks of a given age with PhastCons element overlap.

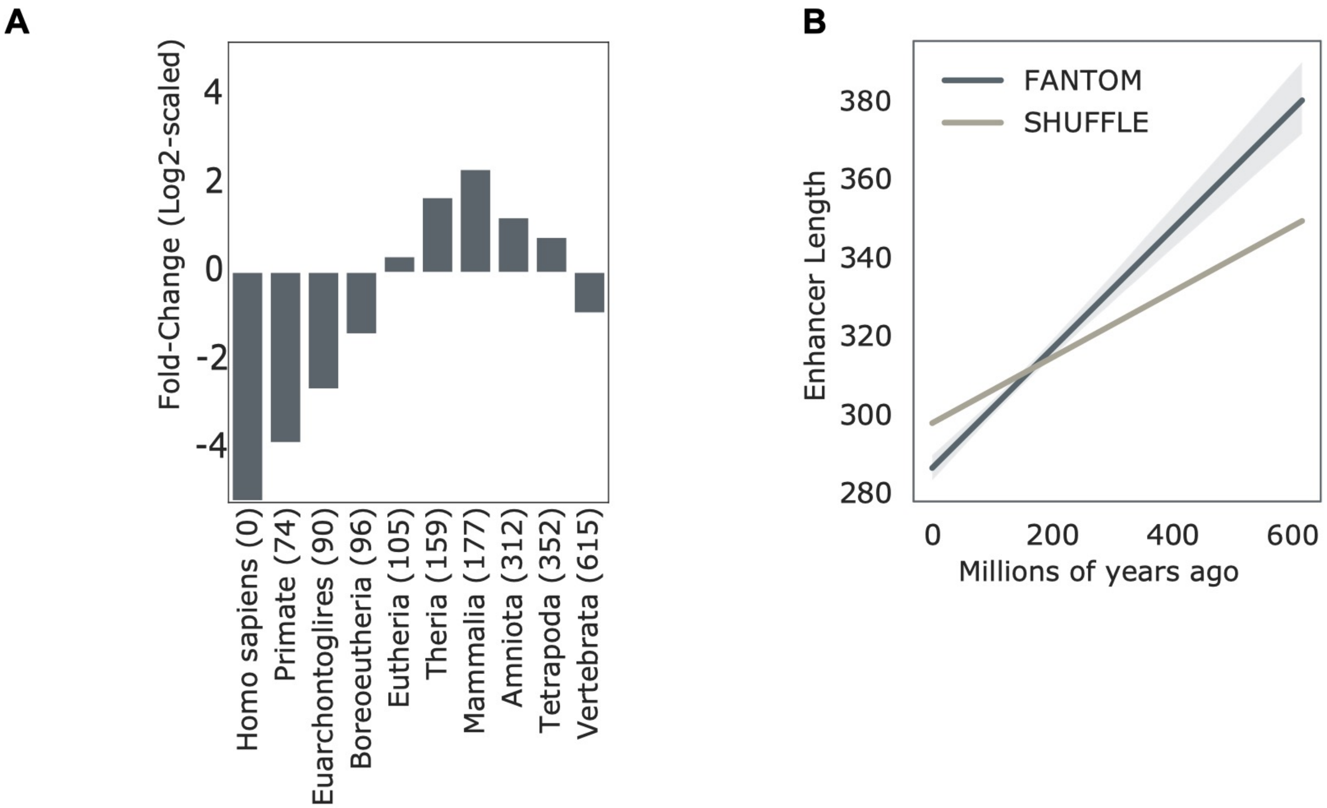


**Supplemental Figure 1.2. FANTOM enhancer sequence ages are enriched for older sequences ages, deplete of younger sequence ages**. (A) FANTOM enhancers are enriched for older sequence ages compared with length-matched random 100x genomic shuffle regions. Fold-change is log2-scaled. Numbers in parenthesis represent estimated MYA since the last common ancestor. (B) Younger FANTOM enhancers are shorter than expectation.

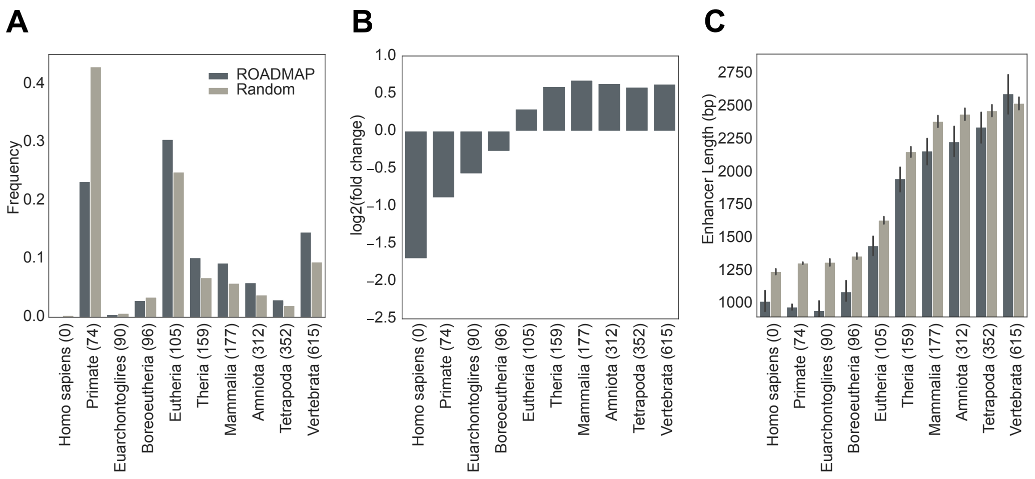


**Supplemental Figure 1.3. ROADMAP Enhancers are enriched for older sequence ages.** (A) Frequency and (B) fold change of ROADMAP enhancer sequence ages (dark grey) across 98 ChIP-seq enhancer datasets versus expected from the genome background (light grey) (mean 0.217 v.0.185 substitutions per site, p =2.4e-39, Mann Whitney U). (C) Enhancer length versus genomic background length per age (p < 2.2e-308, Kruskal-Wallis).

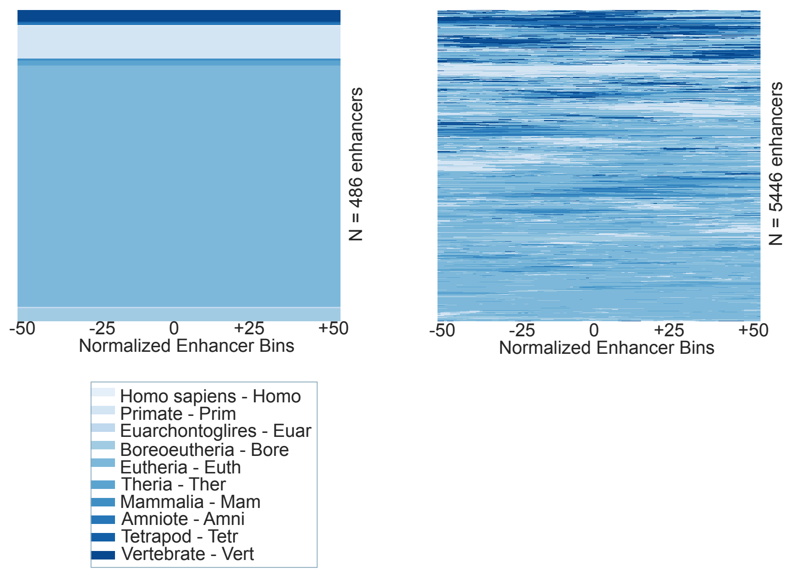


**Supplemental Figure 2.1 – Simple and Complex enhancer age architectures in ROADMAP.** Simple architectures (left) and complex architectures (right) Sequence age architecture sampled from 5932 autosomal ROADMAP enhancers from dataset E072, brain inferior temporal lobe. Enhancer sequence age landscapes were divided into 100 equal-size bins. Age is indicated by color.

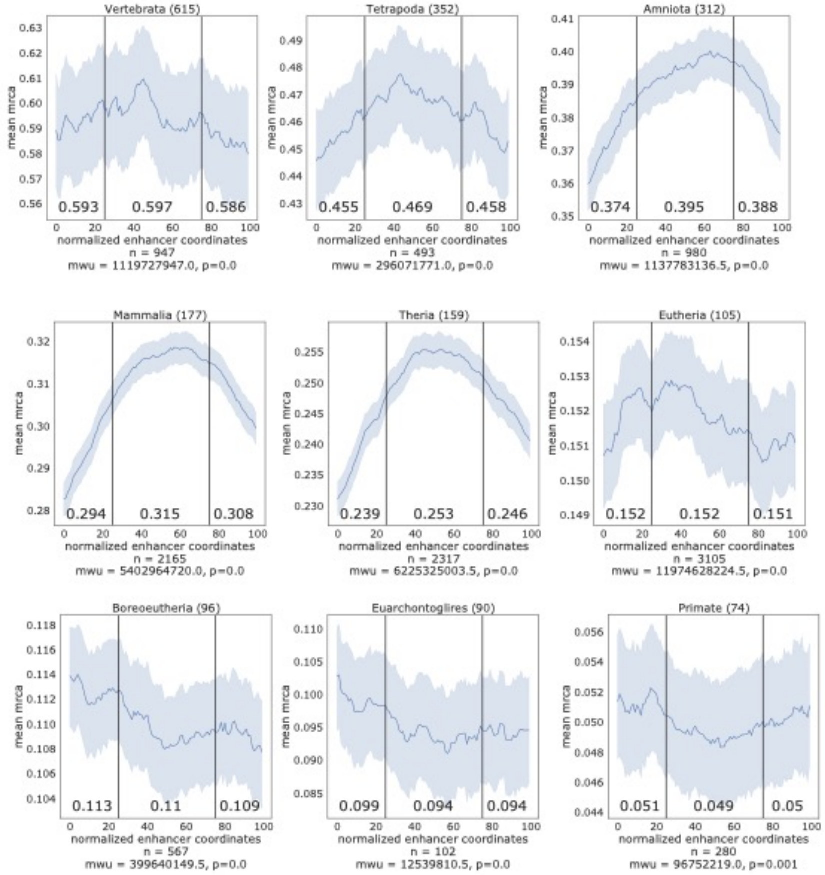


**Supplemental Figure 2.2 –Complex and simple enhancer age architecture landscapes in FANTOM.** Enhancer sequence age landscapes were quantified across 100 bins and stratified by oldest sequence age. Sequence age architecture sampled from 10,956 complex autosomal FANTOM enhancers. Mean age distribution across complex enhancer sequences are shown, one panel per age. Middle 50% versus outer 50% Mann-Whitney U values were calculated for each age classification. Shaded area represents 1000 bootstrapped 95% confidence intervals.

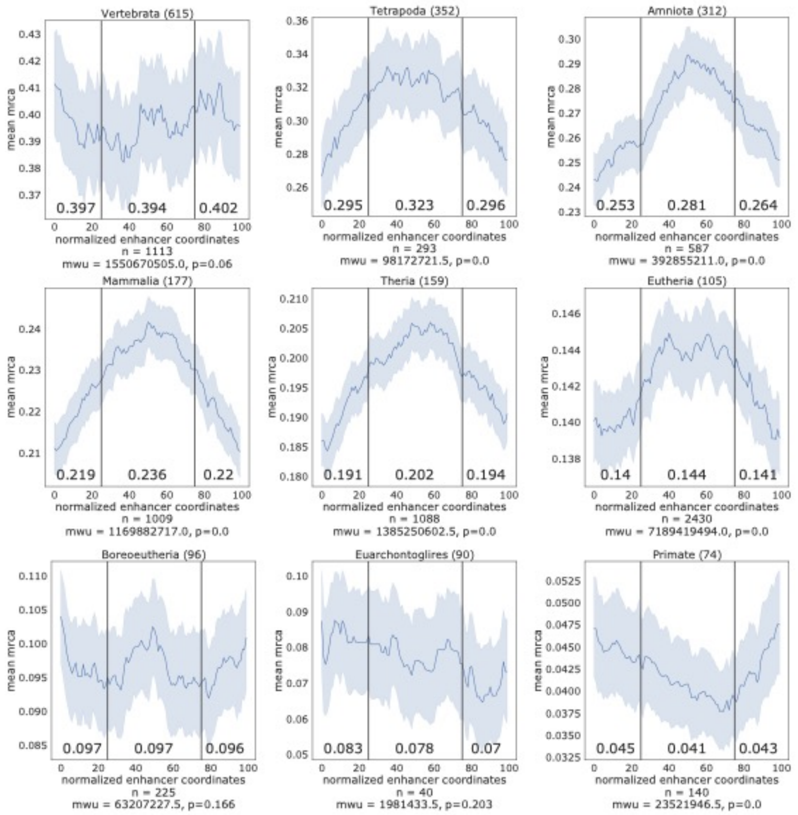


**Supplemental Figure 2.3 Complex enhancer age architecture landscapes in ROADMAP.** Composite landscape of complex sequence age architecture was randomly sampled from 8234 autosomal ROADMAP brain inferior temporal lobe complex enhancers and stratified by age. Enhancer sequence age landscapes were binned into 100 bins and stratified by oldest sequence age. Inner 50% versus outer 50% Mann-Whitney U values were calculated for each age classification. Mean sequence age in substitutions per site for outer quartiles and inner 50% are annotated. Shaded area represents 1000 bootstrapped 95% confidence intervals.

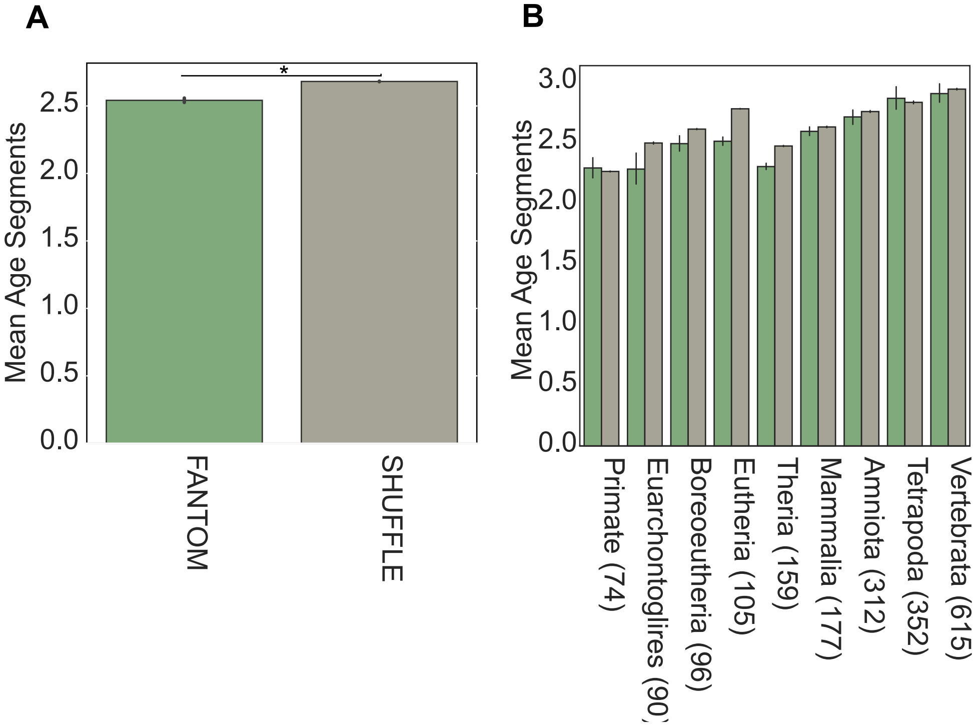


**Supplemental Figure 2.4. Fantom complex enhancers have fewer age segments than expected** based on length-matched regions from the genomic background (overall mean 2.54 complex v. 2.68 shuffle total age segments, $p$ = 9.9e-42, Mann Whitney U). Fantom complex enhancers have significantly different numbers of age segments per MRCA ($p$ = 1.9e-88, Kruskal Wallis) and in comparison with genomic background. Number of segments of different ages in FANTOM complex enhancers of different ages (green) compared to length-matched complex regions selected randomly from the genomic background (gray). At every age, the random segments have greater than or equal numbers of segments of different ages compared to complex enhancers. The largest differences are observed in eutheria and theria enhancers. Error bars are estimated from bootstrapped 95% confidence intervals.

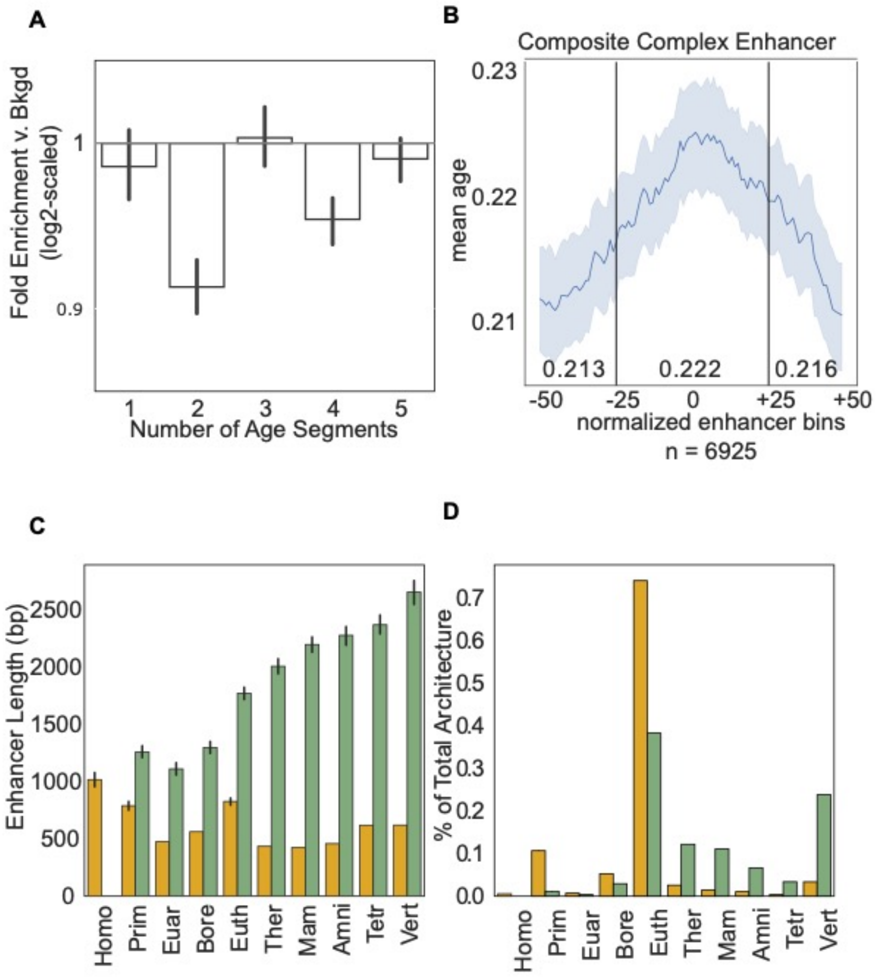


**Supplemental Figure 2.5 – ROADMAP Simple and Complex enhancer age architectures**. (A) Genomic background has more total age segments than ROADMAP enhancers (mean 4.59 v. 4.89 Shuffle v. Roadmap Mann Whitney U, *p* = 0.002). (B) Complex enhancers are oldest at center of enhancer (0.222 inner 50% v. 0.215 outer 50% mean MRCA ages in substitutions per site, *p* <2.2e-308, Mann Whitney U). (C) Complex enhancers are longer than simple enhancers (1843 bp complex v. 542 bp simple median length, *p* <2.2e-308, Mann Whitney U) Enhancer length stratified by age complex and simple enhancers versus genomic background. (D) Complex enhancers are older than simple enhancers, Frequency of complex and simple enhancer architectures per age (overall architecture mean 0.24 v. 0.19 sequence age in substitutions per site, *p* = 2.5e-83, Mann Whitney U)

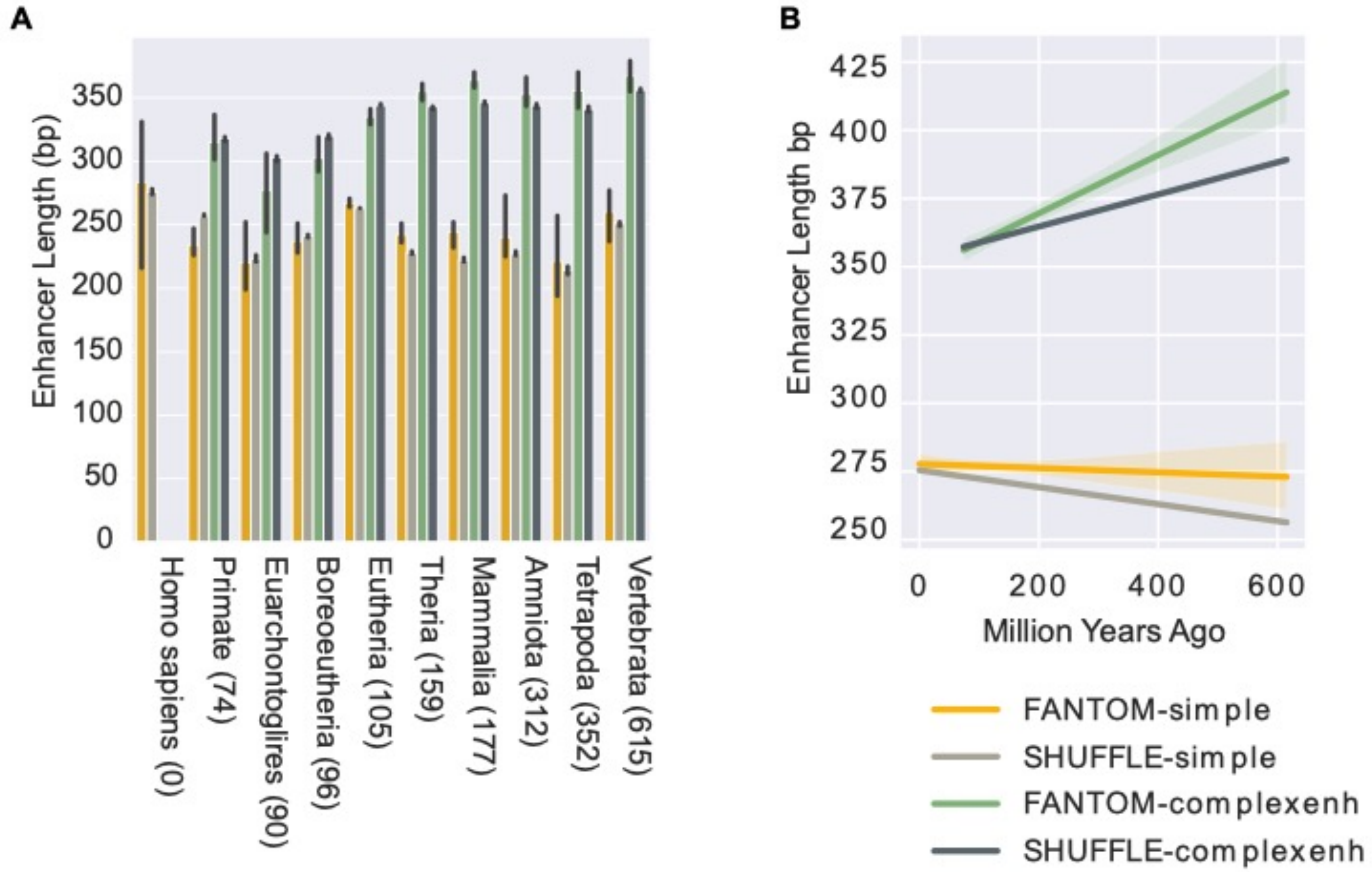


**Supplemental Figure 2.6 – FANTOM simple and complex enhancer architecture lengths versus architecture-matched expectation, stratified by age.** (A) Median ages for FANTOM and shuffled genome architectures stratified by age. (B) Linear regression models of enhancer architecture length per millions of years ago are shown with bootstrapped 95% confidence intervals. Species divergence time estimates were taken from TimeTree (Hedges et al., 2015) . Complex enhancer fitted-line slope = 10.7 bp/100 million years v. complex shuffle fitted-line slope = 5.8 bp/100 million years. Simple enhancer fitted-line slope = -0.8 bp/100 million years v. simple shuffle fitted-line slope = -3.5 bp/100 million years.

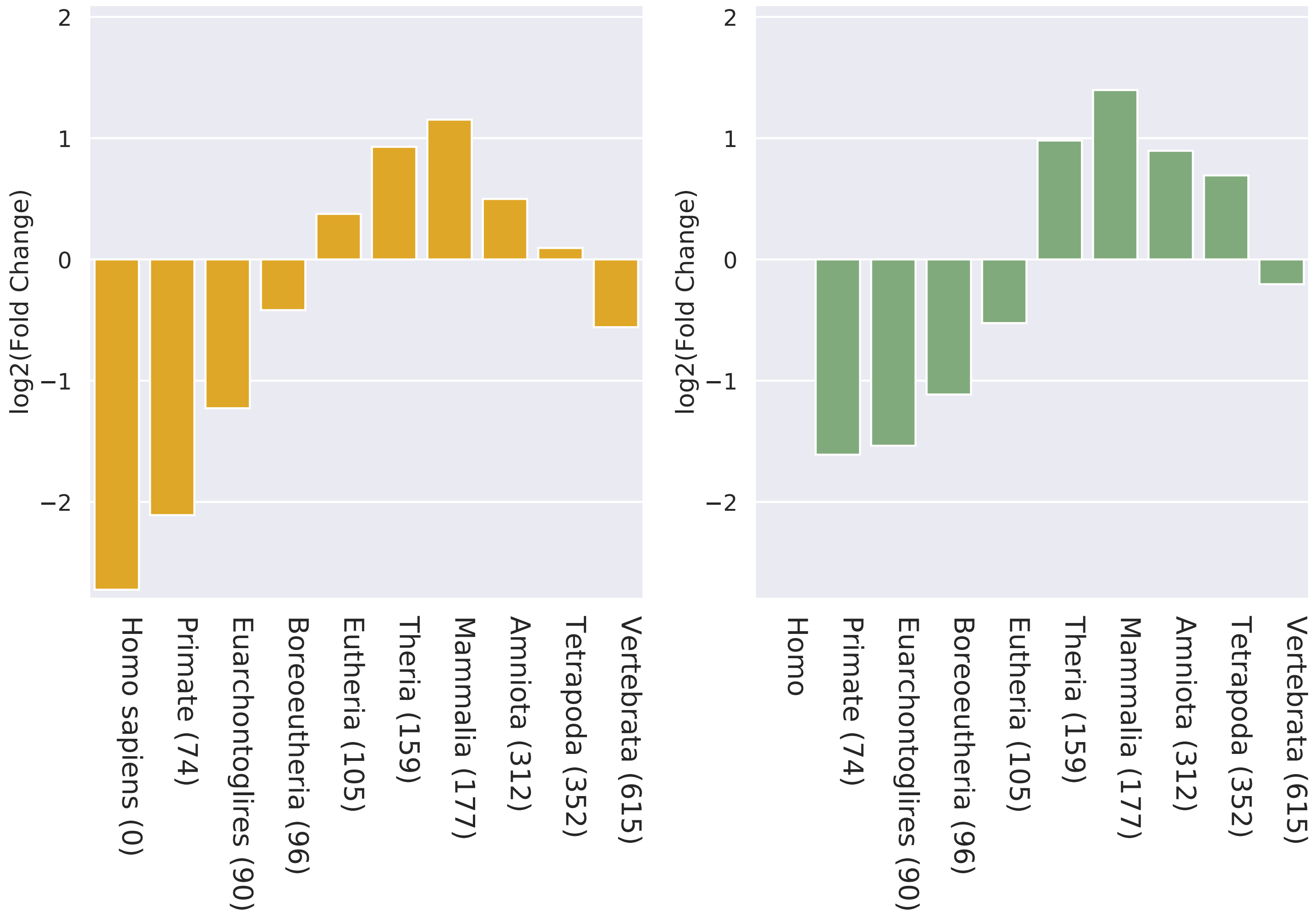


**Supplemental Figure 2.7 – Both simple and complex FANTOM enhancers are deplete of younger sequences, enriched for older sequences compared to expectation**. Fold-change was estimated from simple and complex enhancers against 100x permuted architecture-matched background genome regions.

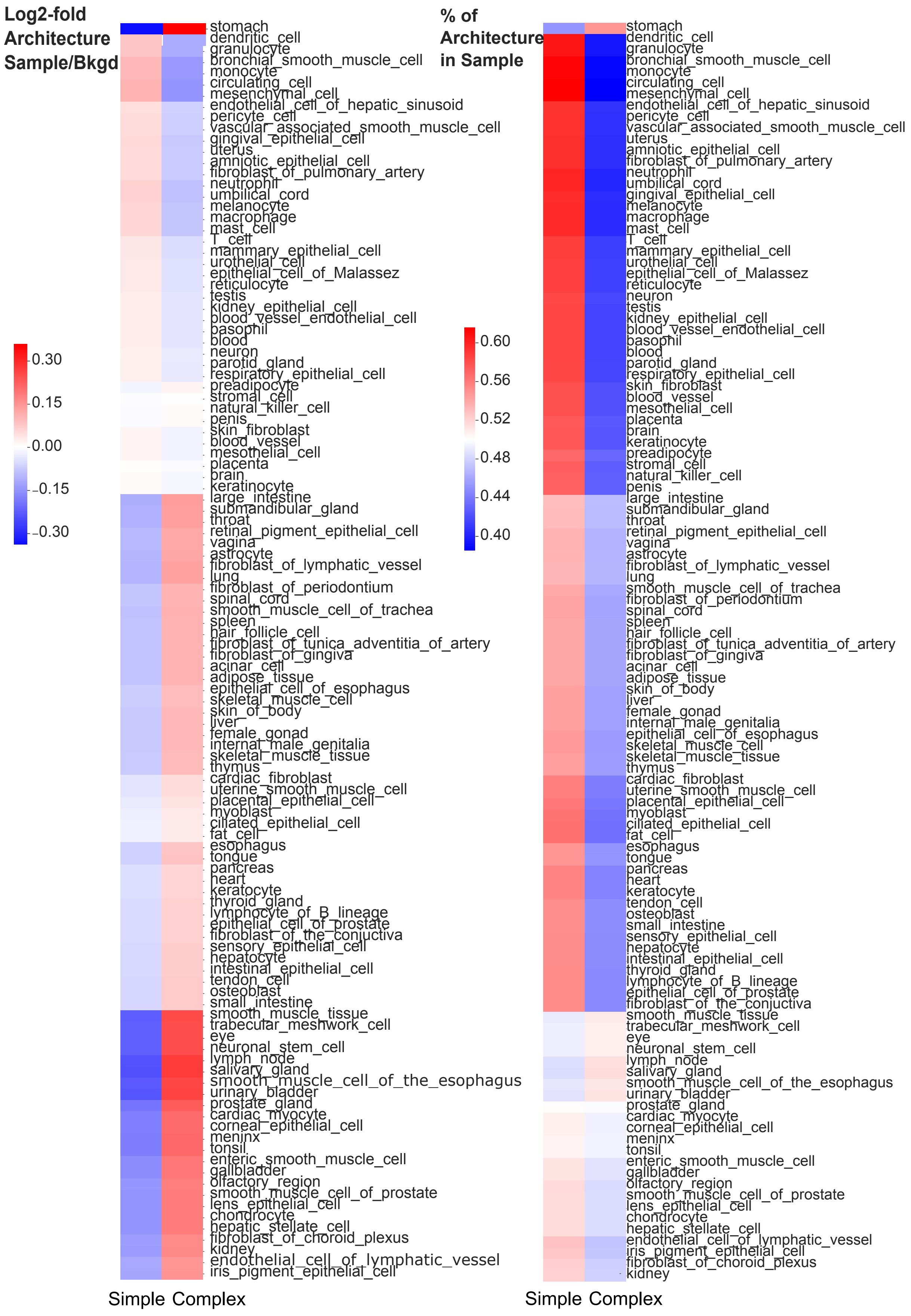


**Supplemental Figure 2.8 – FANTOM simple and complex enhancer architectures per tissue set**. (A) Log2-fold change of simple and complex architectures versus all other tissue datasets sampled in FANTOM. (B) Frequency of simple and complex enhancers across FANTOM cell line and tissue datasets.

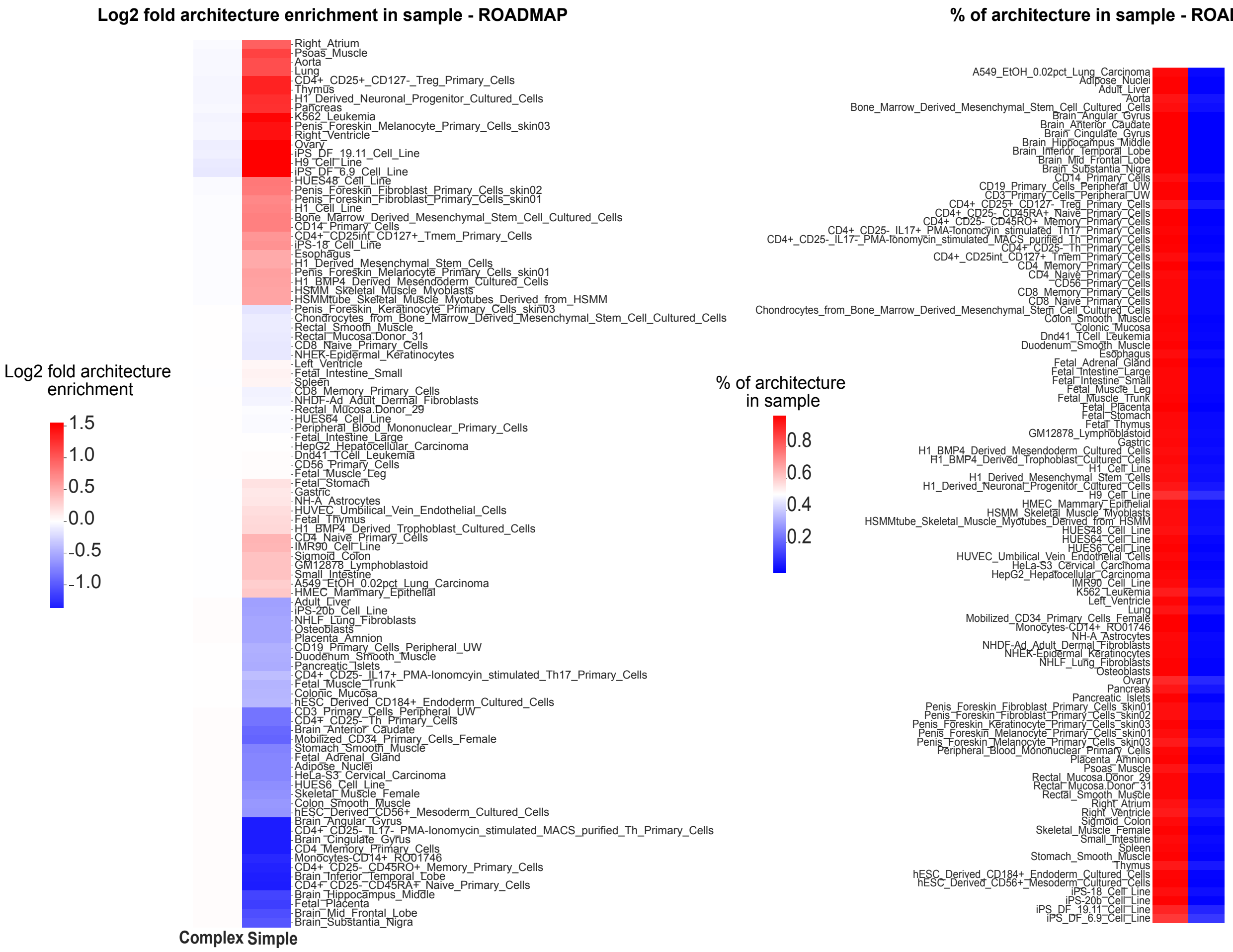


**Supplemental Figure 2.9 – ROADMAP simple and complex enhancer architectures per tissue/cell line dataset**. (Left) Log2 fold change of simple enhancer frequency per dataset versus the median simple enhancer frequency across all datasets. (Right) Frequency of simple and complex enhancers across ROADMAP cell line and tissue datasets.

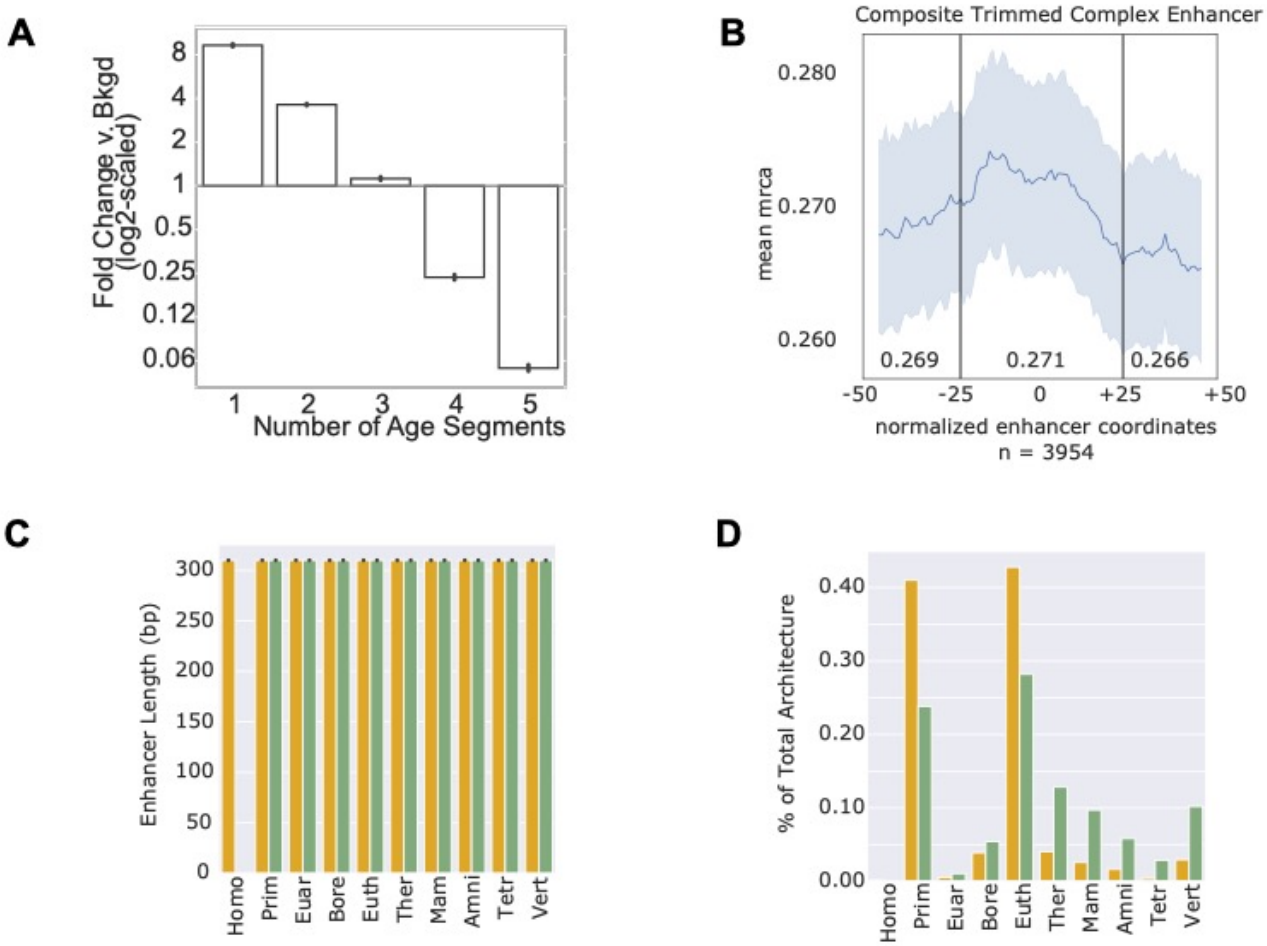


**Supplemental Figure 2.10 – Trimmed ROADMAP Simple and Complex enhancer age architectures**. (A) ROADMAP enhancers are enriched for lower numbers of age segments than expected (odd ratio of 1 age segment = 8.2; *p* < 2.2e-308, Fisher's exact test). (B) Complex enhancers are oldest at center of enhancer (mean 0.222 inner 50% v. 0.215 outer 50% sequence age in substitutions per site, *p* <2.2e-308, Mann Whitney U). (C) Complex enhancer lengths and simple enhancer lengths are equal after trimming, and enhancer length is stratified by age complex and simple enhancers versus genomic background. (D) Complex enhancers are older than simple enhancers, Frequency of complex and simple enhancer architectures per age (overall architecture mean 0.31 v. 0.24 sequence age in substitutions per site, *p* = 8.8e-05, Mann Whitney U)

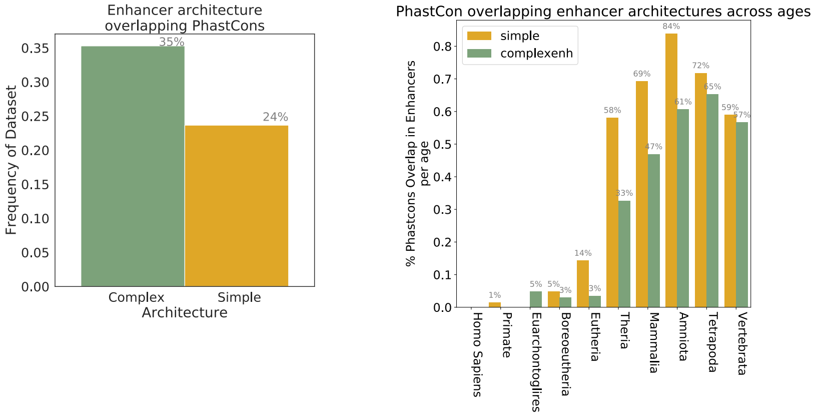


**Supplemental Figure 3.1 – PhastCons estimates for complex and simple FANTOM enhancers.** (A) Complex enhancers are more frequently conserved than simple enhancers. Overall frequency of enhancers overlapping PhastCons elements among simple or complex enhancer datasets. (B) Frequency of enhancers overlapping PhastCons elements within each age.

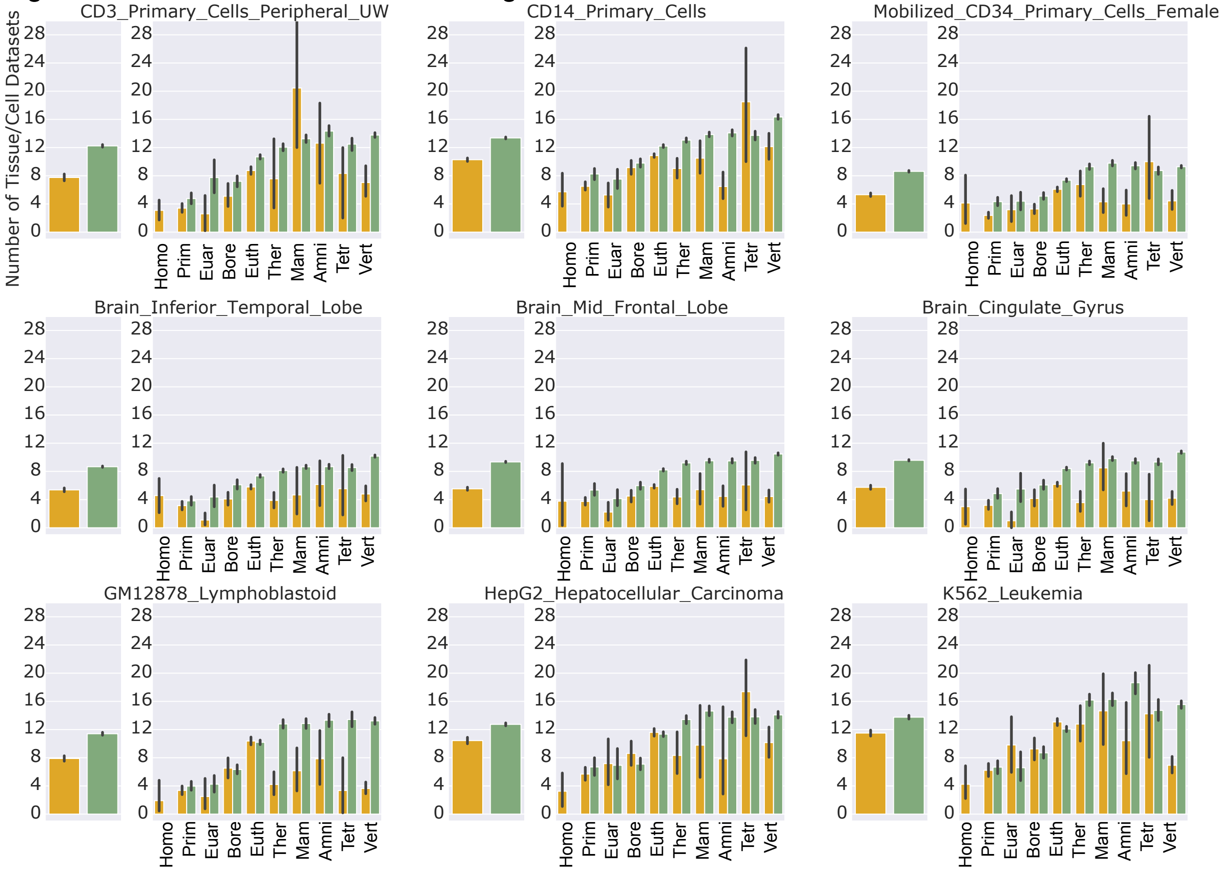


**Supplemental Figure 3.2 – Tissue pleiotropy in simple and complex ROADMAP enhancers.** Roadmap brain, blood, and cell line H3K27ac+ H3K4me3- ChIP-seq datasets were trimmed to mean overall dataset lengths and intersected with 97 other ROADMAP datasets. Mean cross-dataset activities are shown with 95% bootstrapped confidence intervals. Below, cross-dataset activities are shown stratified by enhancer age for each brain, blood, and cell line dataset evaluated.

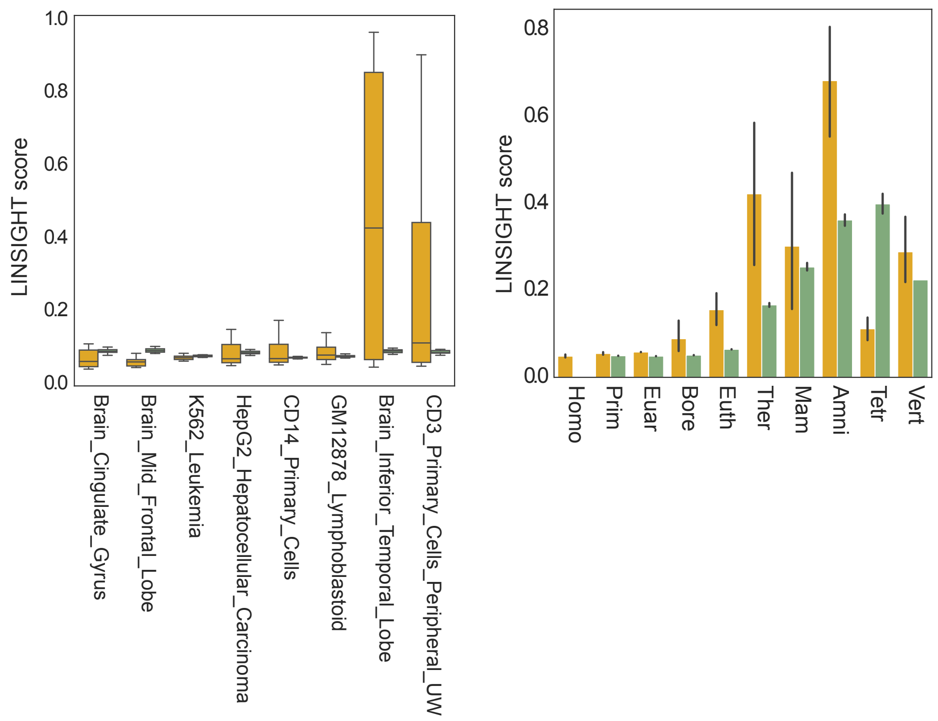


**Supplemental Figure 3.3 - LINSIGHT purifying selection estimates in ROADMAP brain, blood, and cell line H3K27ac+ H3K4me3- ChIP-seq datasets.** (A) Median LINSIGHT estimates for representative ROADMAP brain, blood, and cell line datasets are plotted on the x-axis stratified by architecture. (B) Median LINSIGHT score per architecture and MRCA age combined across ROADMAP brain, blood, and cell line samples.

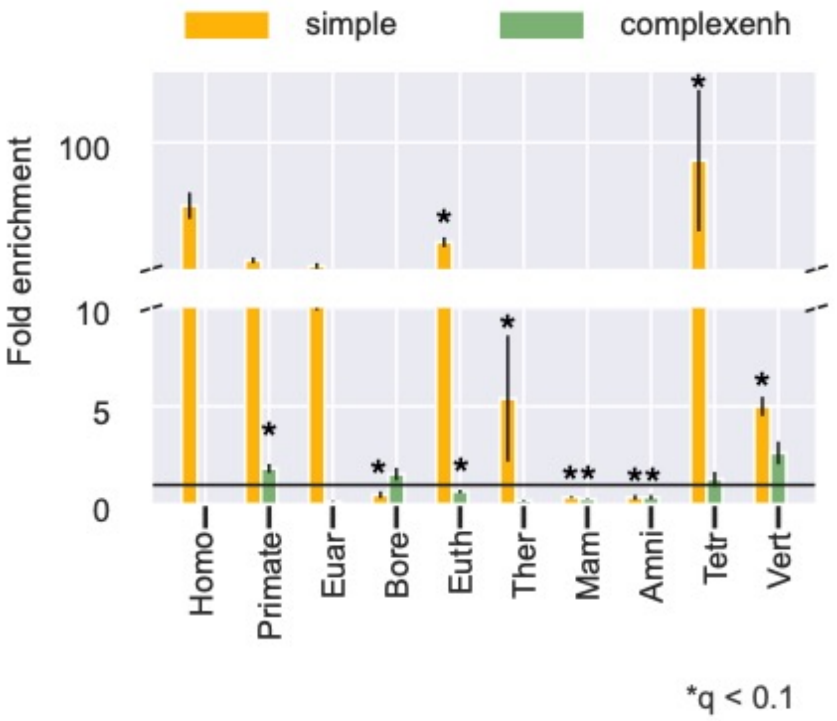


**Supplemental Figure 4.1 – FANTOM simple enhancers are more enriched than complex enhancers for GWAS variants across ages.** Simple and complex FANTOM enhancers were stratified by age and tested for GWAS variant enrichment compared with 100 length-matched and architecture-matched permuted background. Error bars represent 95% confidence intervals bootstrapped 10000 times. Asterisks represent significant enrichment (FDR p <0.1) compared to background.

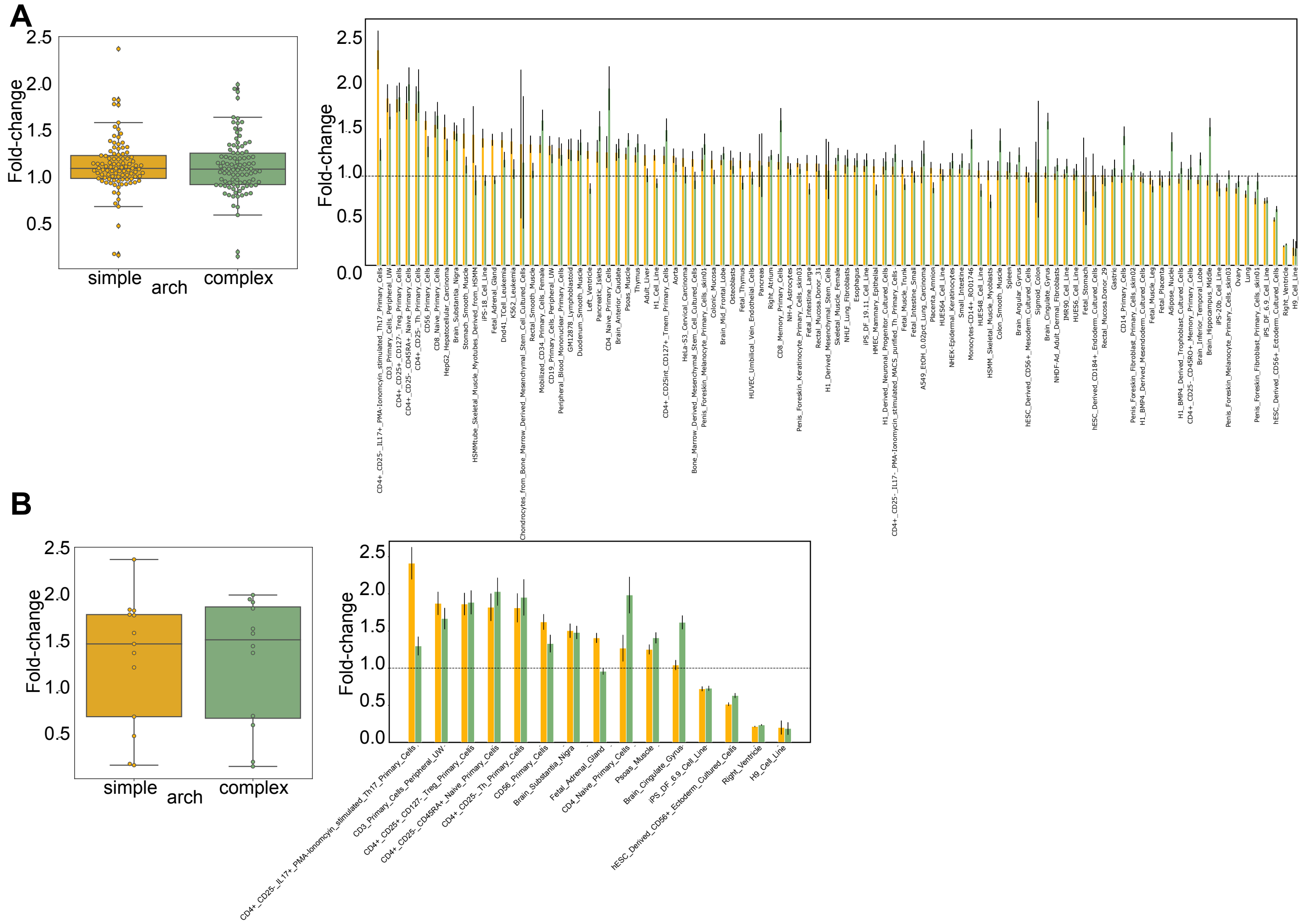


**Supplemental Figure 4.2 – ROADMAP Simple and complex enhancer GWAS tag-SNP enrichment in 97 tissue and cell H3K27ac+ H3K4me3- ChIP-seq datasets.** (A) In 310bp trimmed ROADMAP enhancers among all 97 tissue contexts, simple architectures are more enriched for GWAS variants than complex architectures in 47/97 contexts, while complex enhancers are more enriched for GWAS variants in 48/97 contexts (left panel, median 1.13 simple vs. 1.12 complex fold-enrichment ; $p$ = 0.28, Mann Whitney U). Per dataset comparisons of simple and complex enhancers is shown (right panel). (b) Fifteen tissue contexts showed significant (FDR $p < 0.05$) enrichment of enhancer architecture compared with 100x length-matched, architecture-matched, chromosome-matched background random regions (left panel, median 1.50 simple v. 1.55 complex fold-enrichment; $p$ = 0.38). Among these contexts, 6/15 datasets had higher simple enhancer enrichment, while 9/15 datasets had higher complex enhancer enrichment. Per dataset comparisons of simple and complex enhancers is shown (right panel).

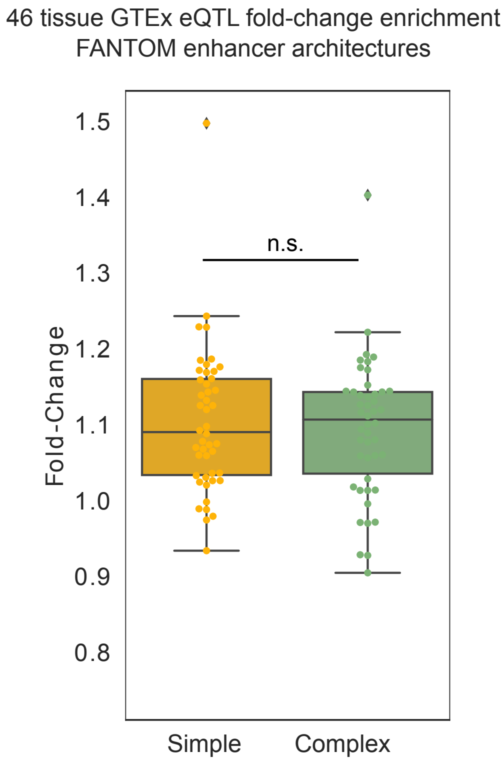


**Supplemental Figure 4.3 – eQTL variants are similarly enriched in simple and complex enhancers** (median 1.09 and 1.11 simple and complex enhancer fold change, $p$ = 0.38, Mann Whitney U). Fold-change enrichment was estimated against a 100x permuted background in 46 eQTL tissue datasets from GTEx. Each dot represents the enhancer architectures eQTL fold-change enrichment per tissue dataset.

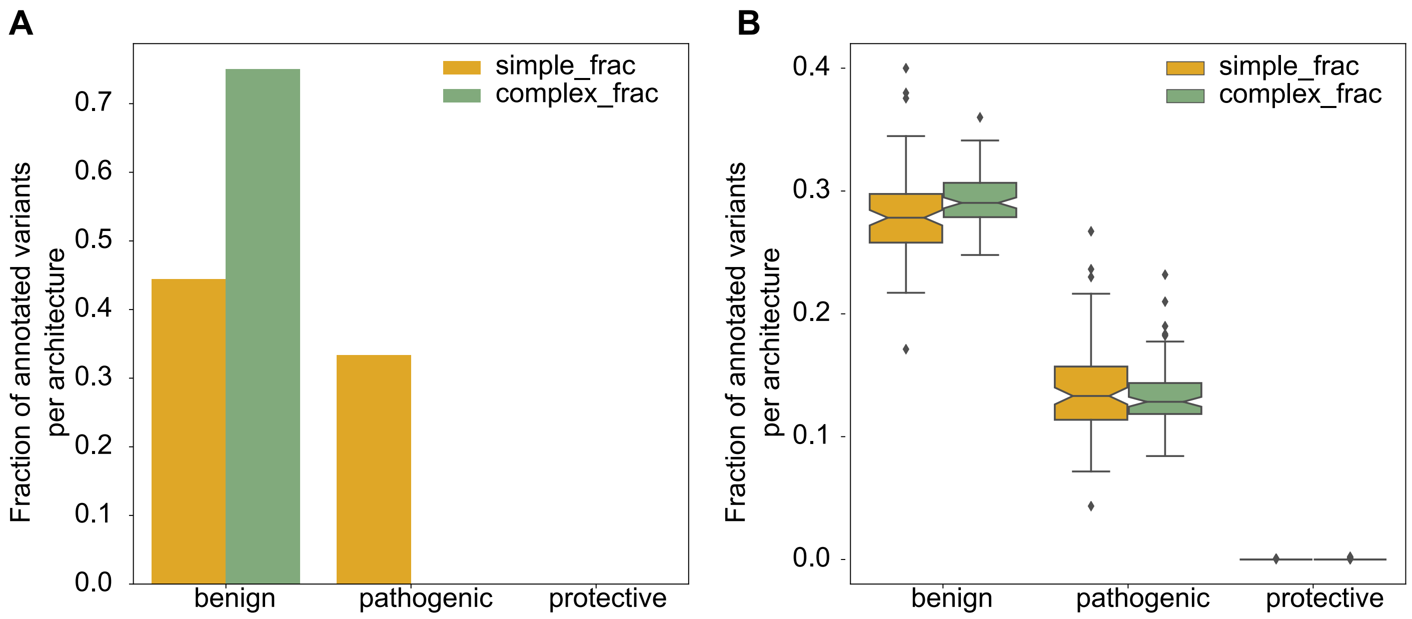


**Supplemental Figure 4.4 – ClinVar annotations in FANTOM and ROADMAP architectures.** (A) Simple FANTOM enhancers are enriched for pathogenic ClinVar variants (0.33 simple v. 0.00 complex enhancer overlaps, $p$ = 0.06 Fisher's Exact Test). Pathogenic annotations include "Pathogenic/Likely_pathogenic" and "Pathogenic_risk_factor". Complex FANTOM enhancers are enriched for benign variants. Benign annotations include "benign" and "Likely_benign". Conflicting annotations were excluded. ClinVar variants were intersected with FANTOM simple and complex enhancers and variant enrichment per annotation was calculated using Fisher's Exact test. Annotations (x-axis) and the fraction of overlapping variants assigned with that annotation (y-axis) are shown. (B) Ninety-eight trimmed 310bp ROADMAP H3K27ac+ H3K4me3- ChIP-seq datasets were intersected with ClinVar variants as done with FANTOM enhancers. Complex ROADMAP enhancers have a higher fraction of benign annotations (median fraction 0.29 complex v. 0.28 simple enhancers, $p$ = 1.17e-4, Mann Whitney U). Simple ROADMAP enhancers have higher fractions of pathogenic variants (median fraction 0.133 simple v. 0.128 complex enhancers, $p$ = 0.28). Neither architecture was enriched for protective variants.

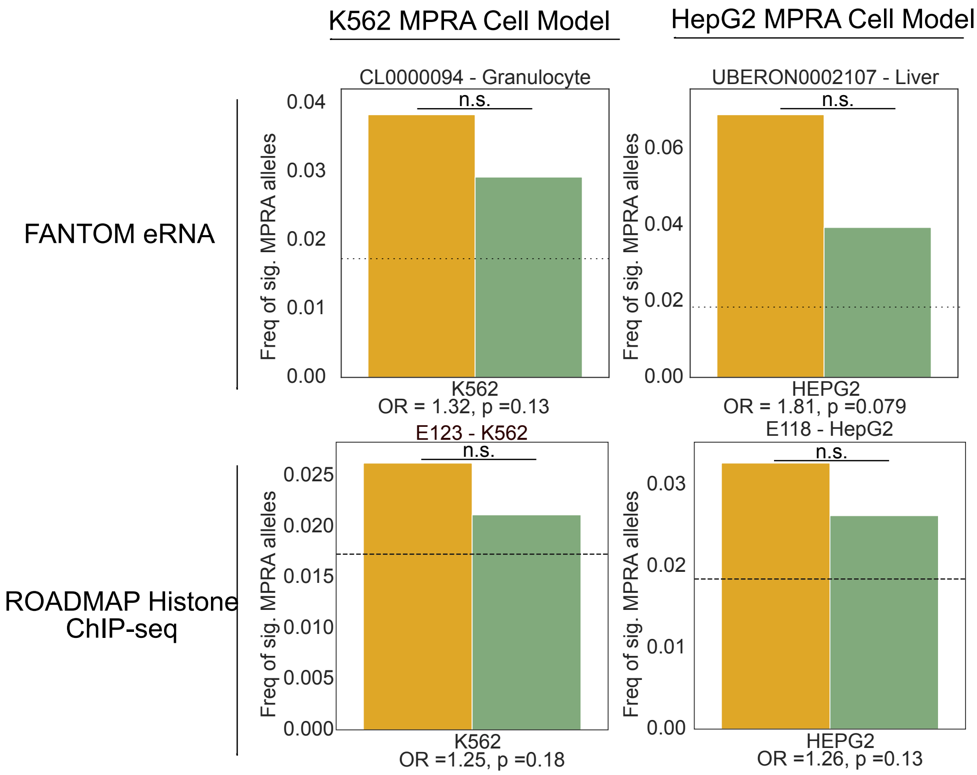


**Supplemental Figure 4.5 – Variants in simple enhancers are enriched for significantly affect regulatory activity in massively parallel reporter assay**. Fraction of tissue/cell-type-specific simple and complex enhancers with significant allelic MPRA activity from FANTOM eRNA and 310 bp trimmed ROADMAP H3K27ac+ H3K4me3- ChIP-seq datasets. Tissue and cell-type-specific datasets were intersected with alleles tested in K562 and HepG2 MPRA assays. The fraction of significant alleles was calculated from all alleles overlapping simple or complex enhancer architectures and is plotted on the y-axis. Enhancer datasets with FDR< 5% significant enrichment are shown in red text. None of the results were statistically significant (*p* <0.05). Significant allelic MPRA activity was determined by the authors using a 5% FDR.

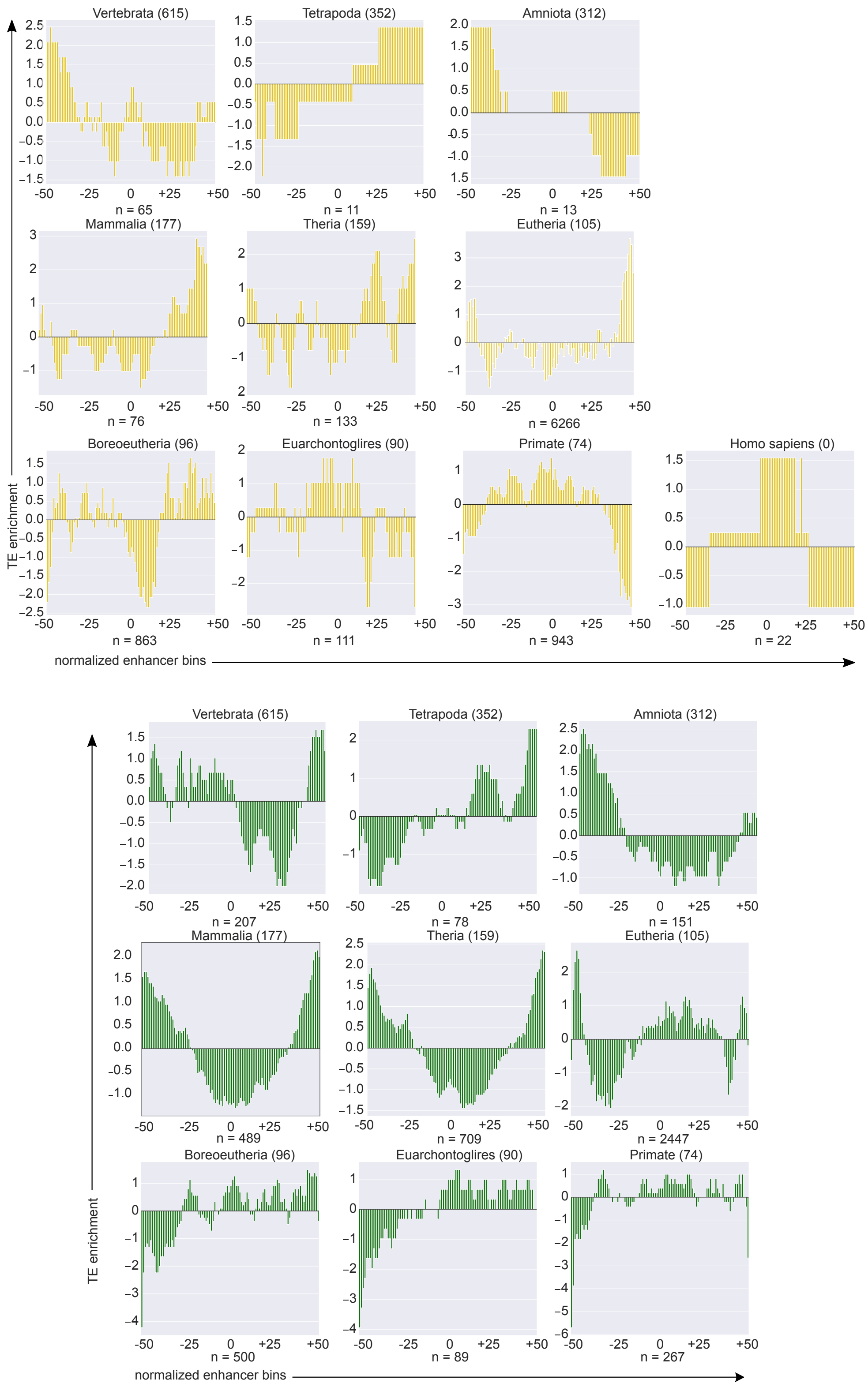


**Supplemental Figure 5.1 – TE enrichment in simple and complex enhancer sequences.** TE enrichment is measured as the z-score of TE overlap counts in normalized enhancer bins and calculated across simple enhancers (yellow) and complex enhancers (green).

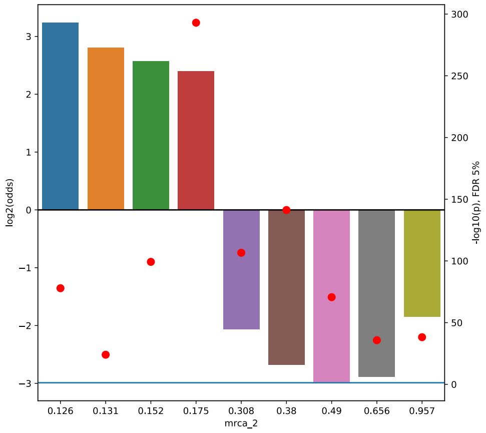


| Complex FANTOM enhancer | TE | No TE |
|---|---|---|
| Core in MRCA | | |
| Core not in MRCA | | |

**Supplemental Figure 5.2 – TEs are enriched in cores of younger complex enhancers, depleted from cores of older complex enhancers.** Log2 odds enrichment of TE overlapping cores in age versus all cores overlapping and not overlapping TEs outside of age. Negative log10(p-value) with a 5% FDR correction is plotted in red dots on the right y-axis.

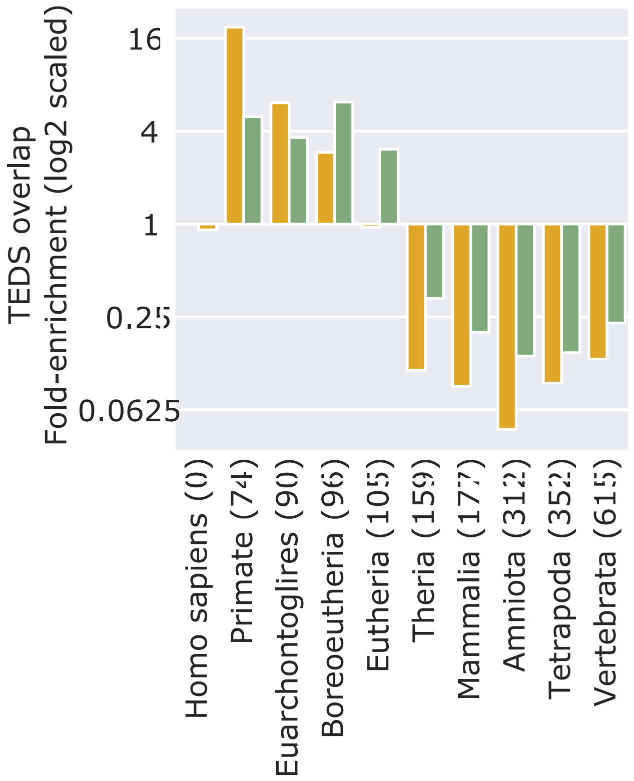


**Supplemental Figure 5.3 – Younger enhancers are enriched for TEs in both architectures**. The ratio of enhancers overlapping TEs to enhancers not overlapping TE within complex or simple architectures per age (median age = 0.175 v. 0.380 for both complex and simple enhancers with TE versus without TE, *p* <2.2e-308, Mann Whitney U).
